# Rv1985c-Driven Transcriptional Network Rewiring Underlies Lineage-Specific Phenotypes in *Mycobacterium tuberculosis*

**DOI:** 10.1101/2020.02.14.943365

**Authors:** Amir Banaei-Esfahani, Sonia Borrell, Andrej Trauner, Sebastian M. Gygli, Tige R. Rustad, Julia Feldmann, Ludovic C. Gillet, Olga T. Schubert, Christian Beisel, David R. Sherman, Ruedi Aebersold, Sebastien Gagneux, Ben C. Collins

**Affiliations:** Department of Biology, Institute of Molecular Systems Biology, ETH Zurich, Zurich, Switzerland; Swiss Tropical and Public Health Institute, Basel, Switzerland; University of Basel, Basel, Switzerland; Department of Global Health, University of Washington, Seattle, Washington, USA; Department of Biosystems Science and Engineering, ETH Zurich, Basel, Switzerland; Faculty of Science, University of Zurich, Zurich, Switzerland; Department of Environmental Microbiology, Eawag – Swiss Federal Institute of Aquatic Science and Technology, Duebendorf, Switzerland; Department of Environmental Systems Science, Institute of Biogeochemistry and Pollutant Dynamics, ETH Zurich, Zurich, Switzerland; School of Biological Sciences, Queen’s University of Belfast, UK

**Keywords:** *Mycobacterium tuberculosis*, infectious diseases, proteomics, genomics, transcriptional network, multi-omics, DosR, Rv1985c

## Abstract

The *Mycobacterium tuberculosis* (Mtb) complex includes ten phylogenetically distinct human-adapted lineages with varying geographical distributions and pathogenicity. Lineage 1 (L1) is associated with low virulence, while Lineage 2 (L2) is linked to hyper-virulence, increased transmission, and drug resistance. We performed multi-layer comparative analyses integrating whole-genome sequencing with quantitative transcriptomic and proteomic profiling of a panel of L1 and L2 clinical strains, each grown under two *in vitro* conditions. Our data revealed variable correlations between transcript and protein levels across strains and gene categories, indicating lineage-specific post-translational regulation. Transcriptional and translational differences scaled with phylogenetic distance, with one in three SNPs on average leading to gene expression changes. A newly developed genome-scale transcriptional regulatory model identified master transcription factors—closely linked to the sigma factor network—whose targets were differentially expressed between L1 and L2. Notably, DosR proteins showed higher basal levels and a stronger nitric oxide (NO) response in L2. Time-course validation using an Rv1985c induction and wild-type H37Rv under hypoxia and reaeration confirmed the role of Rv1985c in the differentially expressed genes between L1 and L2. These findings suggest that limited genetic variation can translate into significant phenotypic differences via differential regulation of key transcriptional networks.

**Highlights:**

- Proteomic and transcriptomic characterization of fully sequenced diverse L1 and L2 clinical strains of Mtb.
- Post translational control mechanisms for regulatory and virulence genes are mitigated in Mtb L2.
- A genome-scale transcriptional framework identifies DosR, Rv1985c, Lsr2 and Rv0691c as master transcription factors responsible for differential target gene expression in L2 compared to L1 strains.
- L1 and L2 DosR proteins respond differently to nitric oxide stress, thus determining a relevant phenotype.
- Time-course phenotypic and transcriptomic data under hypoxia and reaeration validate Rv1985c as a key transcription factor differentiating L2 and L1 strains.

## Introduction

Adaptive evolution is mainly driven by mutations. While the phenotypic consequences of mutations can be easily mapped in the context of strong selection such as antibiotic pressure, the phenotypes emerging from more subtle evolutionary changes are more elusive.

*Mycobacterium tuberculosis* (Mtb) is the etiological agent of tuberculosis (TB), the major cause of human mortality due to an infectious agent ^1^. Mtb is an obligate human pathogen and comprises ten phylogenetic lineages, each with a distinct geographic distribution. Together with several animal-adapted lineages ^2,3^, these make up the *Mycobacterium tuberculosis* Complex (MTBC). Even though the MTBC harbors little genetic diversity compared to other bacteria ^4^, MTBC clinical strains differ significantly in virulence and transmissibility ^5,6^. However, linking this phenotypic variation to the limited genetic diversity in the MTBC has been challenging ^7^. Among the ten human-adapted lineages of the MTBC, some have received particular attention because of their clinical significance. Of note, Lineage 2 (L2) strains are geographically widely distributed and have been associated with high virulence in macrophages and various animal models and rapid disease progression ^8^, increased transmission ^9^, and a propensity for antibiotic resistance ^10^. Conversely, Lineage 1 strains (L1) are geographically localized mainly around the rim of the Indian Ocean, and have been linked to asymptomatic TB^11^, reduced virulence in infection models ^12^, a reduced transmission potential ^9^ and a reduced propensity for antibiotic resistance ^5,13^. Importantly, in geographical areas where both lineages are present, such as in Vietnam, L2 seems to be supplanting L1 as the predominant form of the MTBC ^9,14^. These features point to important physiological differences between L1 and L2. Although, many studies have described phenotypic differences in MTBC clinical strains ^5,15,16^, the molecular mechanisms underlying these differences remain poorly understood. To date, the majority of insights into the molecular processes have been gained from work on a few laboratory-adapted strains of the MTBC ^17–20^ that do not capture the entire phylogenetic diversity of clinical strains^21^.

Microbiologists are increasingly applying systems biology approaches and multi-layer omics technologies to interrogate bacterial systems and their interactions with the human host ^22–25^. These approaches profile various types of biomolecules, typically genome, transcriptome, proteome and, more recently, protein complexes that collectively describe the molecular makeup of specific cells and cellular states and predict interdependencies of different levels of gene expression along the axis of the central dogma ^26–31^. In the MTBC, genomic profiling and measurements of the transcriptional response to a particular stress have been frequently used ^20,32,33^. Yet, computational approaches that can extract mechanistic insights from the statistical associations in a multi-layer omics dataset need further development. Moreover, a growing body of evidence shows that direct functional information can be retrieved from the state of the proteome, making quantitative proteomic data an asset and a relatively recent addition to integrated multi-layered omics analyses of bacteria ^20,34^. The mass spectrometric technique DIA-MS, combines data independent acquisition (DIA) with massively parallel targeted analysis of the acquired data ^35^ and thus offers the high degree of reproducibility, consistency, and throughput needed for large cohort studies. It has been optimized in model experiments with Mtb ^36–38^.

Here, we describe a multi-layer omics dataset that includes transcriptome and proteome data generated from a set of genome sequenced clinical MTBC strains. Specifically, three L1 and L2 strains, respectively, were studied during two *in vitro* growth conditions. We use these data to systematically examine the strength of post transcriptional and translational regulation in the MTBC across various functional categories. Moreover, we tailor a genome scale transcriptional framework – GenSTrans – to identify transcription factors (TFs) that regulate their target genes differently between L1 and L2 strains. We further show that ∼40% of the observed significant changes can be explained by a small set of TFs and the number of SNPs between each pair of strains determines the extent of the significant changes at both mRNA and protein level. Our model highlights four master TFs (DosR, Rv1985c, Rv0691c, and Lsr2) that are known to interact with several sigma factors, including SigB. We discuss how the observed differential expression of DosR proteins, most likely triggered by SigB in response to nitric oxide stress, correlates with shorter growth arrest in L2 strains compared to L1. To validate the biological significance of Rv1985c in differentiating L1 and L2 clinical strains, we design and implement a hypoxia– reaeration time-course experiment in which Rv1985c is induced, thereby supporting the predictive power of our transcriptional model. In summary, we demonstrate the potential of network modeling to describe molecular mechanisms that differentiate MTBC L1 strains from L2. We propose that these differences may explain some of the phenotypic changes observed in clinical settings where these strains circulate.

## Results

### Multi-layer omics profiling of MTBC clinical strains

To establish the data basis for the integrated, multi-layer analysis of MTBC L1 and L2 clinical strains, we selected three drug-susceptible strains from each lineage ^39,40^ to represent the greatest phylogenetic diversity within each lineage based on whole genome sequence data (Fig 1a). We then grew the six strains *in vitro* to mid-log phase and profiled their transcriptomes and proteomes in biological triplicates using RNA sequencing and DIA-MS, respectively (Fig 1b, Fig S1). We quantified 3,933 transcripts across the samples (transcript per million, TPM > 1). DIA-MS quantified 21,184 peptides consistently across samples from which 2,479 proteins, corresponding to 62% of Mtb’s open reading frames, were inferred (Fig 2a, Fig S2a). Through the targeted data analysis pipeline *OpenSWATH* ^41^ and using a Mtb proteome library ^37^ as prior information, three or more proteotypic peptides per protein were quantified for ∼76% of the detected proteins (Fig2b). Next, we examined the quality and reproducibility of each of the omics layer individually. The median Coefficient of Variation (CVs) across the biological triplicates was ∼12% at both the transcript and protein level, indicating comparable data quality between the two platforms (Fig S2b, c). Overall, we were able to quantify 94% of detected proteins across all samples by direct measurement. Where this was not possible, we inferred upper bounds of protein expression from local background values (Fig 2c). The correlation between biomolecular features of the replicates was high for both the transcriptomic and proteomic data (Fig S2 d, e). We assessed the technical and biological variability of the respective datasets. The results indicated an ascending molecular variation with increasing genomic distance within and between the two MTBC lineages (Fig 2d). Our analyses also resulted in a perfect unsupervised clustering of MTBC strains and lineages for both transcriptomic and proteomic data (Fig S2 f, g). Principal component analysis (PCA) suggested that the protein data were more informative with respect to separating strains by their phylogenetic relationship compared to transcriptomic data. Indeed, the first two principal components of protein data exhibited a good separation, resolving individual strains (Fig 2e). By contrast, the resolution of PCA based on transcriptomic data was more limited and reflected the phylogenetic classification (Fig 2f). In short, we have generated a dataset, which spans from the genome of fully sequenced clinical isolates of MTBC to transcriptome and proteome. The multifaceted nature and deep coverage of the high data quality provided a solid base on which to build our downstream analyses.

**Figure 1.**
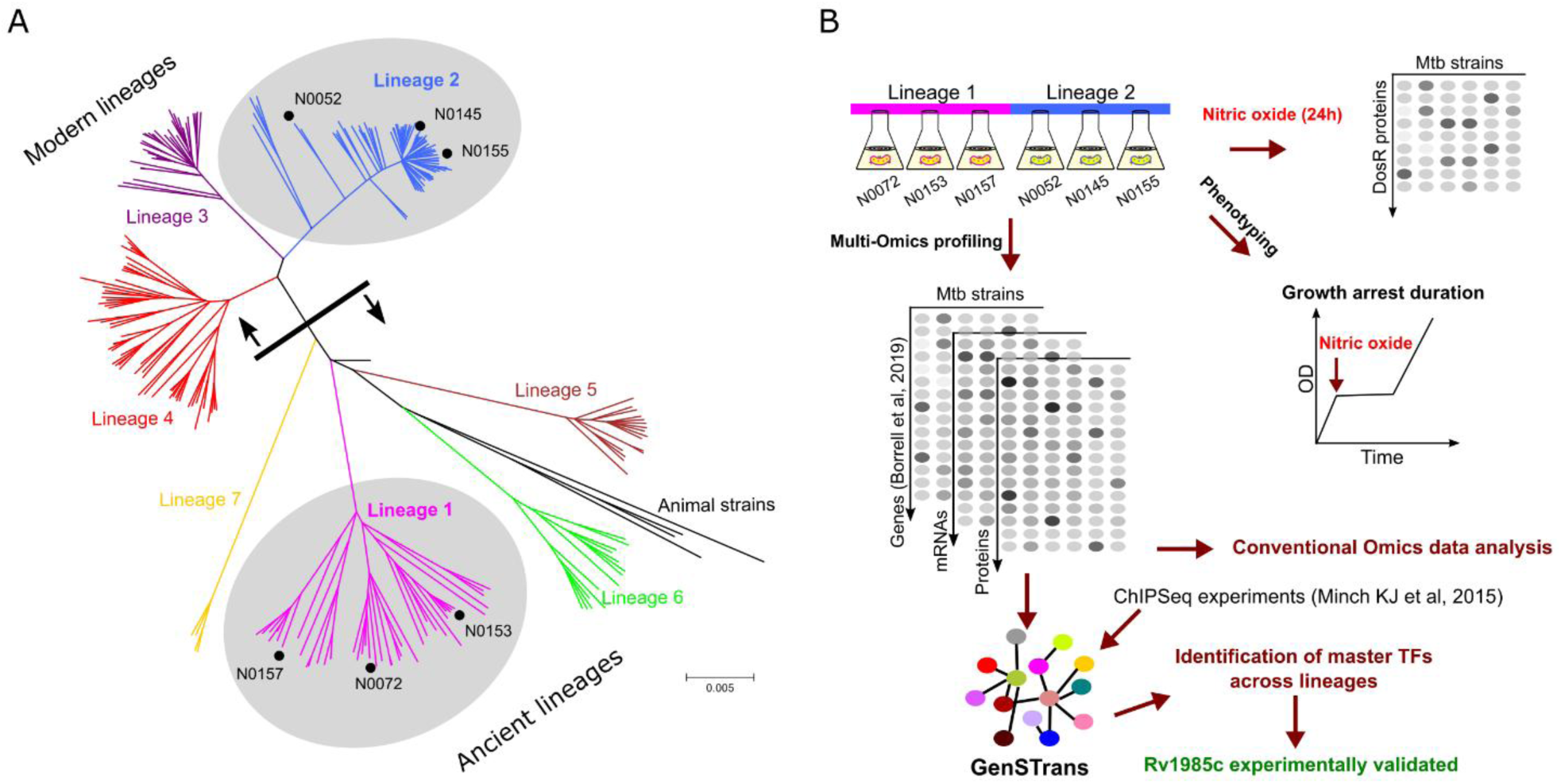
Schematic illustration of workflow. **(A)** Dendrogram shows the evolutionary relationships between the seven human-adapted lineages of the *Mycobacterium tuberculosis* complex and the L1 and L2 strains that were selected for this study. **(B)** Conceptual workflow. Transcriptome and proteome of six fully sequenced clinical strains belonging to L1 and L2 were measured. A genome-scale transcriptional model was developed to identify transcription factors regulating their targets differently between the two lineages. An exemplary phenotypic consequence of the differentially regulated DosR proteins between the two lineages and in response to a stress condition, the growth arrest duration following nitric oxide exposure.

**Figure 2.**
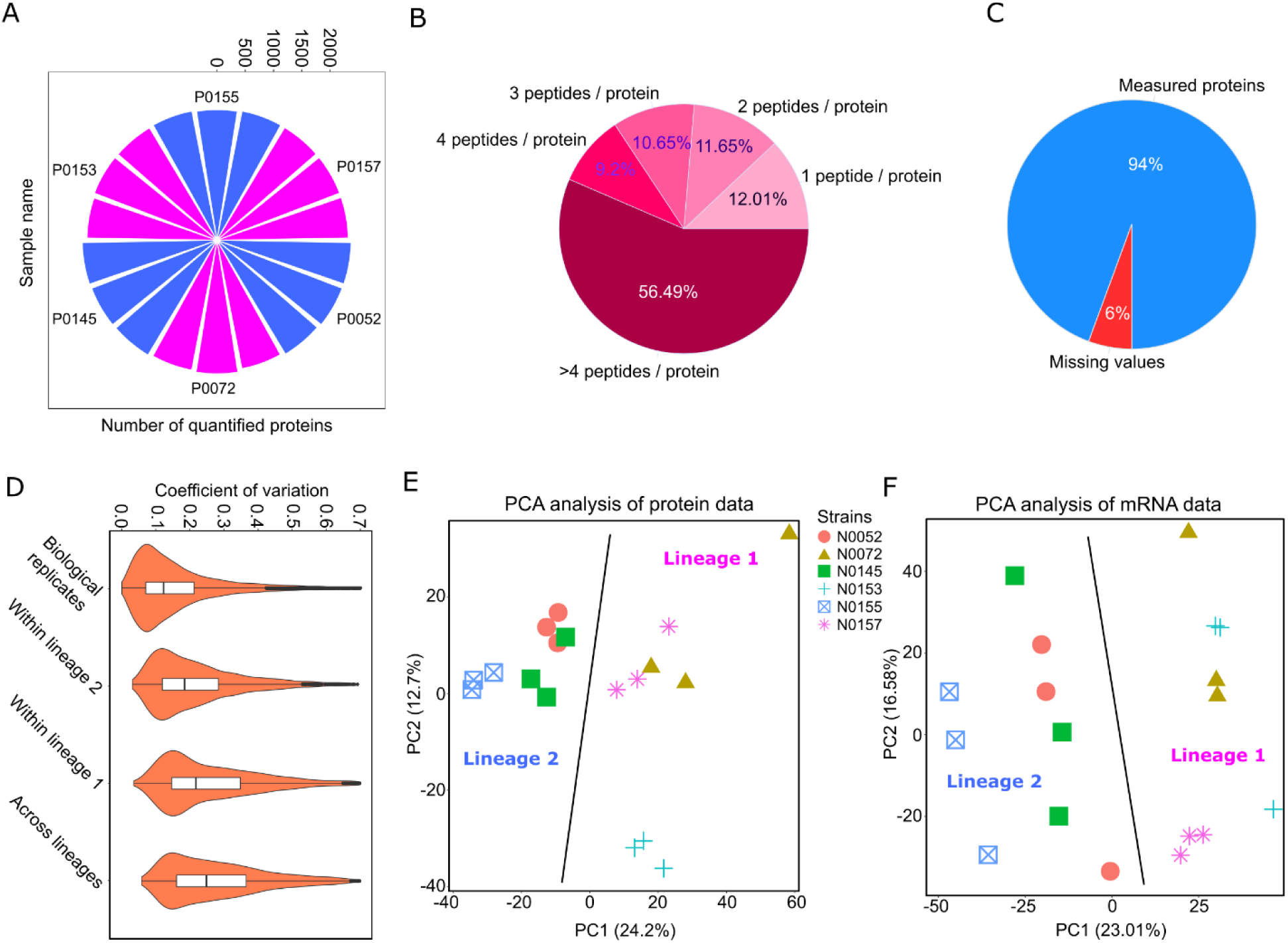
Cellular protein landscape in MTBC clinical strains. **(A)** Number of proteins quantified in each biological replicate of a clinical strain at 1% protein FDR. **(B)** Pie chart exhibits the deep coverage of peptides identified by DIA-MS at 1% protein FDR. **(C)** Completeness of protein measurements. 94% of proteins across the samples were quantified by direct peptide measurements while the remaining (6%) were inferred using background signals. **(D)** Variation in protein measurements due to different factors: technical (median CVs = 12%), within L2 (median CVs = 19%), within L1 (median CVs = 23%) and across all measurements (median CVs = 26%). **(D)** PCA analysis of protein data. PCA plot shows a decent separation resolution at the strain level. **(E)** PCA analysis of mRNA data. PCA analysis of mRNA data could separate clinical strains based on only their phylogenetic lineages.

### Post-translational regulation is prevalent in MTBC clinical strains

We used our combined multi-layer omics dataset to systematically explore the extent of correlation between mRNA and protein levels in MTBC clinical strains and to assess the relative impact across gene functional categories annotated by Mycobrowser ^42^ between the two lineages. We computed the Spearman rank correlation between averaged intensities of mRNA and protein products of the same gene across biological triplicates in the six clinical strains studied. We found that the magnitude of the correlation, and hence the assumed extent of post transcriptional and translational regulation, varied between functional categories. The overall correlation was 0.46, slightly higher than in previous reports ^20^. The strongest and weakest signals of post transcriptional and translational regulation were observed in the “*virulence, detoxification, and adaptation*” (Spearman’s rho: 0.33) and “*lipid metabolism*” (Spearman’s rho: 0.69) group, respectively. Remarkably, the L2 strains presented a significantly higher mRNA to protein correlation compared to L1 for both regulatory (T-Test’s p-value = 0.015) and virulence genes (T-Test’s p-value = 0.0036) (Fig 3a). A growing body of evidence has revealed that the extent of post-transcriptional regulation in the MTBC is largely limited, as indicated by the high correlation (r = 0.93) observed between mRNAs and ribosome-protected fragments (RPFs) ^19^. This suggests that the modest correlation between mRNA and proteins may instead originate from post-translational regulation in the MTBC. Comparing the global correlation observed in the MTBC to that in other species revealed that the MTBC’s potential for post-translational regulation could significantly decouple its proteome from its transcriptome, to a degree comparable to that observed in humans, despite the generally lower complexity of bacterial systems. ^23,43^. An earlier study showed that mRNAs and proteins of 1,921 *Escherichia coli* genes were largely correlated (r = 0.8), supporting the notion that post transcriptional and translational regulation are relatively limited in bacteria ^44^. Other studies in yeast and metazoan species showed mRNA-protein correlations at the gene-to-gene basis of above 0.5 ^45–48^. A possible explanation for the apparent increased post translational regulation in the MTBC may be the presence of the Pup-proteasome system responsible for protein degradation and homeostasis that is rarely found in other bacteria including *E. coli* ^49^. Due to the substantial post translational regulation in the MTBC, RNA data in isolation can lead to incomplete or misleading mechanistic understanding. This underscores the need for high quality quantitative proteome data ^20^. For instance, our data revealed that T cell antigens (both MHC class one and two) are significantly regulated at the RNA level in L2 strains compared to L1 (MCHI’s p-value = 4.57E-5 and MHCII’s p-value = 0.0071). However, this pattern of the differential regulation could not be observed at the protein level (MHC I: p-value = 0.07 and MHC II: p-value = 0.55) (Fig S3).

**Figure 3.**
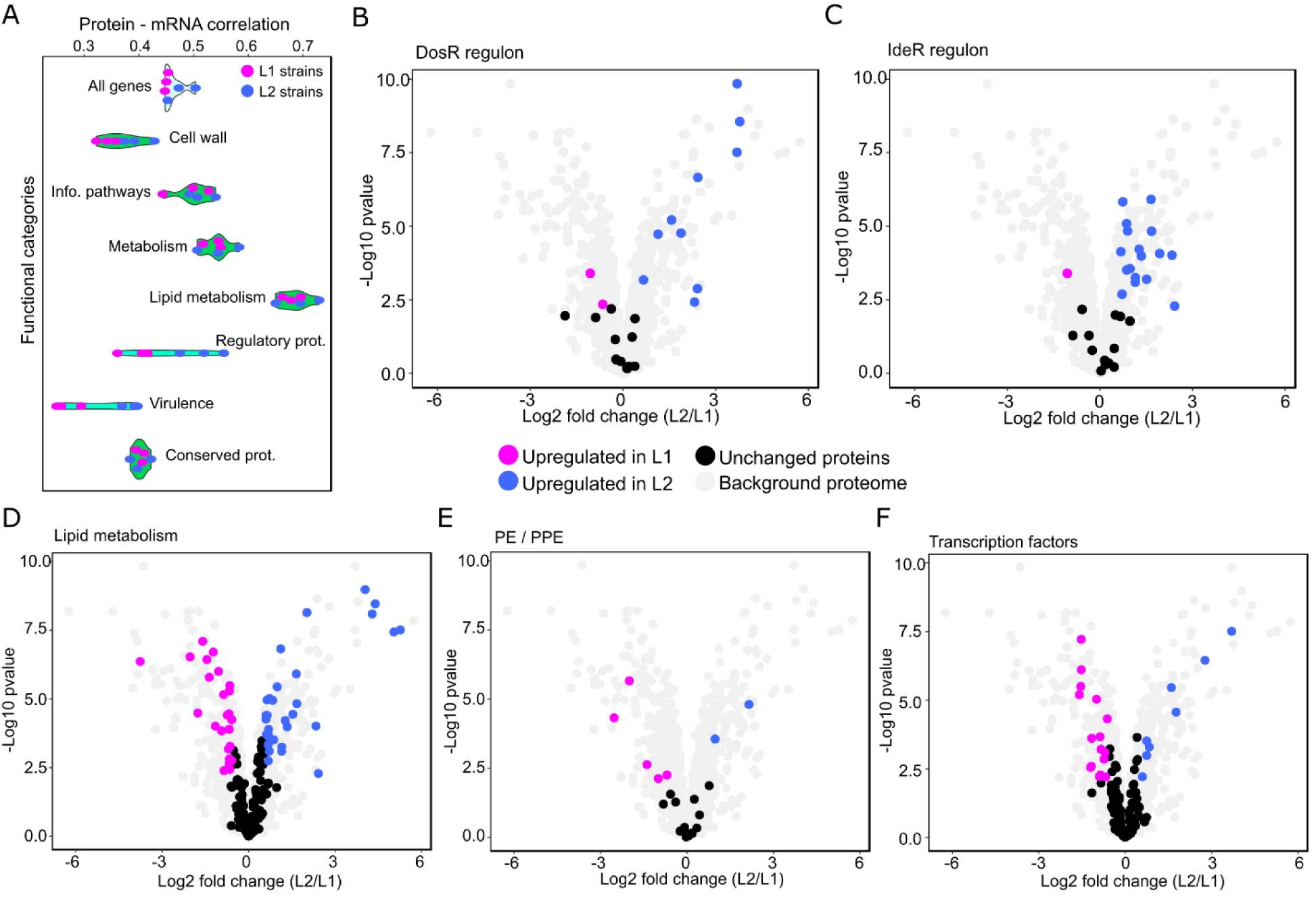
Enrichment and correlation analyses of MTBC functional categories. **(A)** Spearman correlation analyses of protein vs. mRNA abundances at the gene-to-gene basis for different functional categories: all genes (average R = 0.46), cell wall and cell processes (average R = 0.37), information pathways (average R = 0.50), Intermediary metabolism and respiration (average R = 0.54), lipid metabolism (average R = 0.69), regulatory genes (average R = 0.46), virulence, detoxification, and adaptation genes (average R = 0.33) and conserved genes (average R = 0.40). Volcano plots showing proteins of different functional groups that are differentially expressed between L1 and L2: **(B)** Dormancy survival Regulator (DosR) regulon (p-value = 2.5E-5), **(C)** Iron-dependent Repressor (IdeR) regulon (p-value = 8.3E-9), **(D)** lipid metabolism (p-value = 3.6E-5) and **(E)** PE/PPE genes (p-value = 0.014). **(F)** 24 transcription factors (out of 138 quantified) showed significant changes in their abundance in L2 strains compared to L1. For (B) to (F), purple and blue dots denote proteins significantly (p-value < 0.01 and fold change > 1.5) upregulated in L1 and L2 strains respectively, and black dots denoting proteins of the respective groups that do not significantly change. Figure S4 contains the corresponding mRNA data.

### Transcript and protein abundances differ substantially between MTBC L1 and L2 strains

Next, we performed differential protein and transcript expression analyses of the set of L1 and L2 strains. We identified 578 differentially abundant transcripts and 390 differentially abundant proteins (fold change > 1.5 and adjusted p-value < 0.01) (Table S1). The vast majority of the changes occurred in non-essential genes involved in survival, infection and persistence of the bacilli, whereas the expression of essential genes remained largely unchanged (p-values = 6.9E-7 and 2.17E-6 on mRNA and protein level, respectively). Gene set enrichment analyses of the differentially abundant transcripts and proteins highlighted various regulons and functional categories that distinguish the two lineages. These included the Dormancy survival Regulator (DosR) and Iron-dependent Repressor (IdeR) regulons. DosR proteins contribute to maintenance of cell viability during transition to a non-replicating state induced by various stresses such as macrophage engulfment, hypoxia and nitric oxide ^50^. The abundance of these genes was significantly differently regulated in L2 strains compared to L1, both on the transcript and protein level (Fig 3b, S4a). For transcripts this observation has been reported previously ^26,51,52^ and it is confirmed here using both transcriptome and proteome data from the same samples. Chao JD *et. al.* showed that the abundance of DosR proteins is increased upon knockout of the serine/threonine-protein kinase PknH ^53^. Since we found the abundance of PknH decreased in L2 strains, we decided to examine whether there was a link between the PknH downregulation and DosR upregulation. We hypothesized that PknH could be involved in the differential regulation of DosR proteins, in addition to the causative variant C601T ^54^, through either DosR or an alternative transcription factor. To test this hypothesis, we profiled the proteome of a PknH knockout and overexpressing strain generated from the laboratory strain H37Rv, along with the wild type strain in biological triplicates, using the same method as described above. However, we could not replicate the results shown by Yossef Av-Gay and colleagues and DosR proteins remained unchanged upon PknH knockout and overexpression (Table S1).

Our analyses also pointed to the regulon IdeR, (repressed in iron-replete conditions) as being significantly differentially expressed between the two lineages (Fig 3c, S4b). Iron acquisition systems in pathogenic bacteria are critical virulence factors and the MTBC requires iron to be able to grow in culture and during infection ^55,56^. Hence, the increased expression of IdeR proteins in L2 strains could boost the potential of the iron acquisition system and therefore strengthen L2’s success during infection ^57^. Lipid metabolism related proteins that showed the highest mRNA-protein correlation also appeared significantly differentially abundant between the two lineages (Fig 3d, S4c). This functional category contains 272 genes in Mtb whereas, in comparison, the *E. coli* genome encodes only ∼50 such proteins. The amount of energy the MTBC spends to transcribe and translate these genes indicates their critical roles in the life cycle of the MTBC. Proline-glutamic acid (PE)/proline-proline-glutamic acid (PPE) genes, many of which are considered as virulence factors, also displayed differential regulation between the lineages at both RNA and protein level (Fig 3e, S4d). These proteins are challenging to quantify since they are, to a large extent, either secreted or localized on the cell surface. We next examined transcription factors (TFs) of which 138 out of 214 (64%) known factors could be detected at the protein level. We found that at least 24 TFs showed changes in their abundance at the protein level between the two lineages (Fig 3f, S4e). Taken together, our multi-omics data revealed important differences in the functional and regulatory organization of MTBC clinical isolates from L1 and L2.

### The number of single nucleotide polymorphisms (SNPs) is linked to the magnitude of transcript and protein expression differences between strains

The large extent of the observed transcriptional and translational changes between the two lineages led us to examine the role of SNPs, the most common form of genetic variation in the MTBC. To this end, we first calculated the genomic distance based on number of SNPs between each pair of the tested strains. The average pairwise genomic distances within L1 and L2, as well as between the two lineages, were 873, 446 and 1,761 SNPs, respectively. These results are consistent with results reported previously ^5^ that showed that the genetic diversity within L1 was about twice that of L2. We then examined the extent to which the observed changes at the transcript and protein level correlated with the number of SNPs in the corresponding genomes. Our analysis revealed a Spearman correlation between the genomic distance, measured in number of SNPs, and the differentially regulated mRNAs and proteins of 0.73 and 0.83, respectively (Fig 4a, b). The lower proteomic coverage diminished the slope of the corresponding regression line but it almost met the slope calculated for transcripts following a coverage-based normalization. The elucidation of the one-to-one relationship and exact mechanism of each causative SNP goes beyond the scope of this analysis due to the complexity of potential mechanisms. For instance, genetic variation might affect regulatory proteins such as TFs by either changing the affinity of a given protein to DNA (both synonymous and non-synonymous SNPs), or modifying a regulatory protein (non-synonymous SNPs), and therefore influence gene expression. Hence, we included both synonymous and non-synonymous SNPs in our analysis. Overall, our data indicate that, on average, every three SNPs (1/slope of regression line) caused a significant expression change which can be detected at either the transcript or protein level or both.

**Figure 4.**
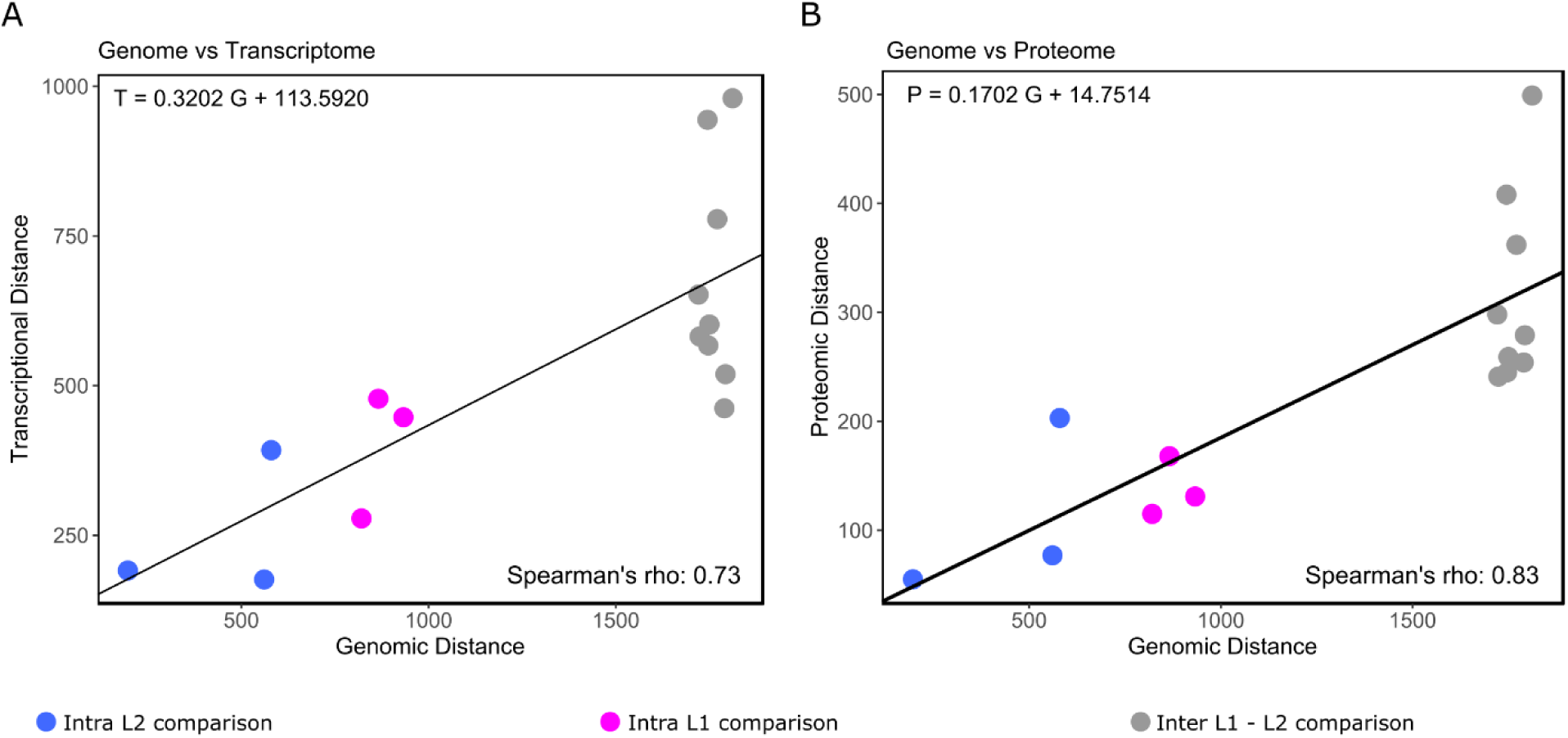
Genetic distance determines number of significant changes on mRNA and protein level. **(A)** transcriptomic distance and **(B)** proteomic distance, measured by number of significant changes on mRNA and protein level respectively, as a function of genetic distance, calculated based on number of distinct SNPs for a given pair of clinical strains. Their slopes (following coverage-based normalization for proteomic slope) and high correlations showed that every third SNPs, on average, causes a significant change at the transcript and protein level.

### A genome-scale transcriptional model identifies divergent regulators between L1 and L2 strains

The differential expression of a high number of TFs between the two lineages observed in our dataset led us to ask to what extent changes in transcript or protein abundance can be explained by direct TF regulation. To address this question, we leveraged an existing transcriptional framework, the Mtb’s Environment and Gene Regulatory Influence Network (EGRIN) model, to identify TFs causally linked to the observed changes in transcript and protein abundance ^58^. The EGRIN model defines clusters of co-regulated transcripts (modules) and their corresponding TFs. It requires dual enrichment analysis to connect a given regulated gene set to the modules and the modules to TFs (Fig S5a) ^59^. Applying the model to our transcriptomic dataset highlighted 15 modules that were i) associated with at least one TF and ii) significantly enriched in transcripts that were differentially abundant between the two lineages (adjusted p-value < 0.05) (Fig S6a). The transcription factor DosR was identified with high significance through four different modules as a putatively causative TF for divergent gene expression in L1 and L2 strains (Fig S6b). In addition, we also found that KstR2 (Rv3557c) could regulate its target genes differently between the two lineages. This transcription factor was identified through two different modules and controls the expression of a small set of genes involved in the utilization of cholesterol, the main carbon source used by the MTBC during infection ^60^. Whereas EGRIN provides a high degree of sensitivity and is well suited to pathway-level analysis provided that the model includes pathways of interest, the model could not satisfactorily address our quest for the identification of lineage-specific regulators for two reasons. First, the EGRIN model contains only 66 transcription factors and is therefore not a genome-scale model. Second, each module encompasses only 13 genes on average, which mitigates the specificity of the model. Consequently, the Mtb EGRIN based analysis can suggest several TFs regulating a given module, making it impractical to identify the causative TF with confidence (Fig S6a). To improve on these limitations, we devised a new genome-scale transcriptional framework.

We repeated the analyses with a new genome-scale transcriptional network model, GenSTrans, that was developed based on ChIPSeq experiments ^61^. The network consists of 143 transcription factors and their corresponding sets of target genes represented as specific sub-networks. Each sub-network on average consists of 64 genes. The high density of information in the network is expected to boost the specificity of the analysis compared to the EGRIN model. We overlaid the transcript data of this study on the GenSTrans network and examined the degree of enrichment of differentially abundant transcripts in specific sub-networks using hypergeometric test followed by Benjamini-Hochberg based multiple testing correction (Fig S5b). This analysis suggested 28 TFs that differentially regulated gene expression between L1 and L2 strains (Fig 5a). Of the 28 TFs identified by GenSTrans, 17 were not identified by the EGRIN model due to insufficient genome coverage (Fig 5a). Our new model could recapitulate at least one TF corresponding to 11 modules also identified by the EGRIN model (Fig S6a). Of note, GeneSTrans more precisely pointed to one or two TFs per EGRIN’s module in cases where the EGRIN model has suggested several equally likely candidates. For instance, the EGRIN model identified module 446 that could potentially be regulated by either Rv0081 or Rv1460, or a combination of both (Fig S6a). Rv0081 and Rv1460 bind to 647 and 11 genes, respectively, according to ChIPSeq experiments^61^. Deconvoluting the signals is further complicated because the module overlaps with both sub-networks significantly. Nevertheless, the non-overlapping genes pointed to Rv1460 as the most likely causative TF explaining the observed pattern, as 9 out of 11 genes in the corresponding sub-network appeared regulated in the comparison of L1 and L2 strains.

**Figure 5.**
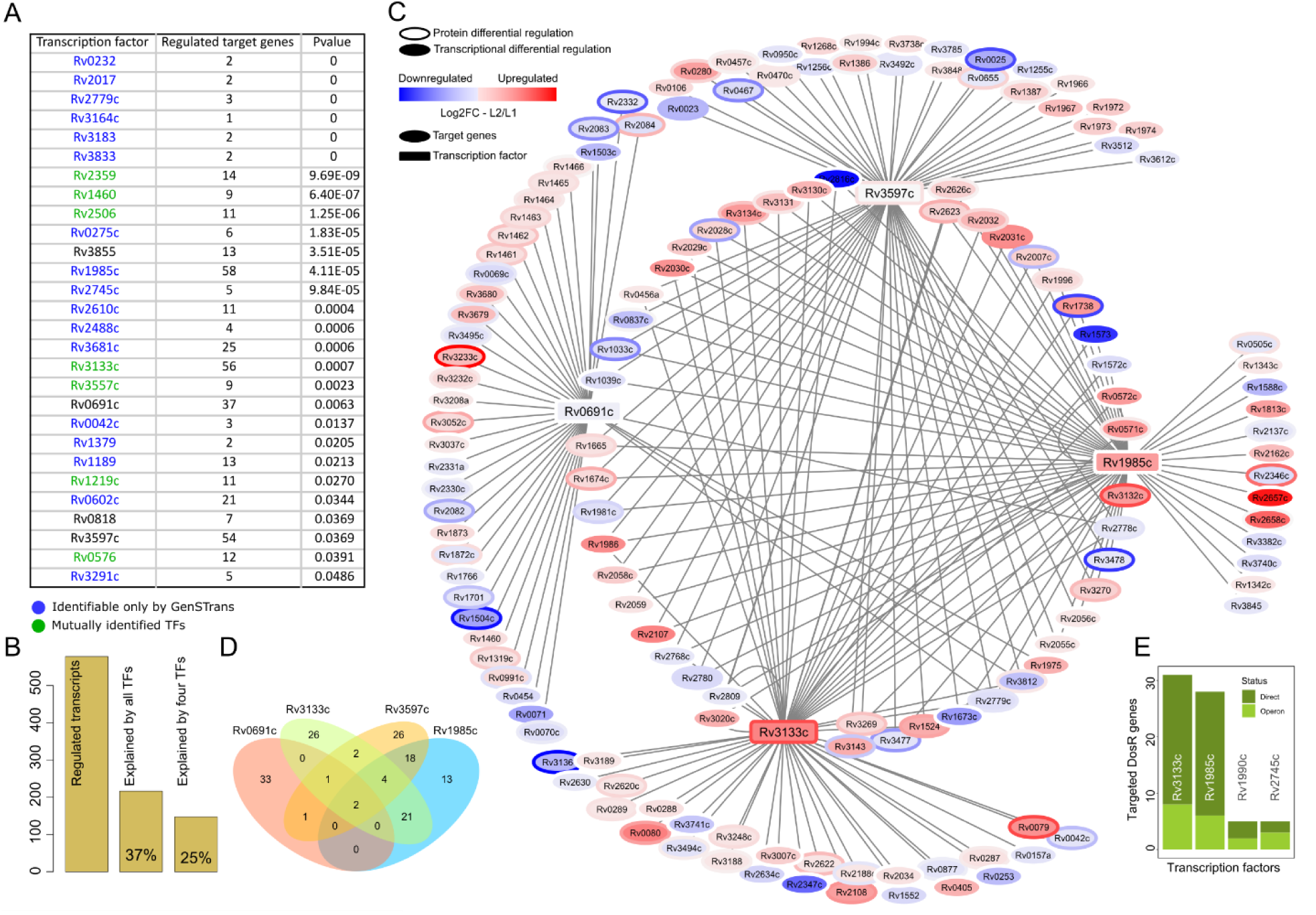
Transcriptional modeling in MTBC L2 strains compared to L1. **(A)** List of 28 transcription factors regulating their target genes differently between L1 and L2 strains. Blue gene names denotes TFs that are not identifiable by the EGRIN model due to its sparse coverage. **(B)** Subnetwork of four master transcription factors explaining 25% of mRNA changes in L2 strains compared to L1. Node and border colors depict transcript and protein expression respectively. **(C)** Bar plot summarizes percentiles of regulated transcripts between the two lineages that could be explained by identified and four master transcription factors. **(D)** Venn diagram of target genes that are regulated by four master transcription factors. High degree of overlapping genes particularly between Rv3133c (DosR) and Rv1985c suggested their close functional relationships. **(E)** Bar plot summarizes number of DosR Regolun genes regulated by various TFs. It further confirms that Rv3133c (DosR) and Rv1985c work closely together.

Next, we returned to the original question about causation of differential gene expression between L1 and L2 strains. Specifically, we examined the extent to which the identified TFs could describe the observed expression differences. Our results revealed that at least 37% of the modulated transcripts could be statistically significantly linked to a relatively small set of TFs. A set of only four TFs – Rv3133c (DosR), Rv1985c, Rv3597c (Lsr2), and Rv0691c – were sufficient to explain a quarter of the transcriptomic changes between L1 and L2 strains (Fig 5b). DosR on both transcript and protein level and Rv1985c on transcript level – not identified through our proteome profiling – were differentially regulated in their abundance very significantly between the two lineages. Interestingly, and in contrast to DosR and Rv1985c, Lsr2 and Rv0691c did not show significantly different transcript/protein abundance between the lineages. We speculated that they might undergo post-translational modifications (PTMs) regulation that affect their respective level of activity. In the MTBC, various types of PTMs () have been identified on at least 2,617 proteins ^30,62^. For instance, PknB, a serine/threonine-protein kinase, can phosphorylate Rv3597c ^63^. The sub-networks of these four TFs shared several genes with each other, indicating their close functional relationship (Fig 5c, d). Of note, DosR and Rv1985c regulate 31 and 28 DosR regulon genes, respectively (Fig 5e), according to ChIP-Seq experiments, with a significant gap between them and other TFs in terms of their involvement in regulating DosR regulon genes ^61^ (Fig S7). These highly overlapping target genes belonging to the DosR regulon might indicate Rv1985c as an alternative TF contributing to the DosR target regulation under certain conditions.

### Hierarchical organization for interactions between TFs and sigma factors

In addition to 214 TFs, the tubercle bacillus has one housekeeping sigma factor, SigA, and 12 accessory sigma factors leading to the reprogramming of RNA polymerase (RNAP) and consequently the initiation of transcription of particular gene sets. The sigma subunit ensures specificity of the RNAP holoenzyme for special promoter sequences ^64,65^. Gennaro and colleagues reconstructed a sigma factor regulatory network of Mtb. It consists of 13 sigma factors, seven anti-sigma factors, two anti-anti-sigma factors and 50 TFs ^66^, altogether 72 genes. Of these, we were able to quantify the abundance of all 72 genes at the transcript level and 48 genes at the protein level, allowing us to ask how significantly the four master TFs could interact with the sigma factor network and generate differential responses to a given stress.

We assessed whether the four master TFs identified in our analysis were present in the sigma factor network and how their interactions with sigma factors/anti-factors might affect gene expression (Fig S8). Three of the four master TFs, Lsr2, DosR and Rv0691c, were among the 50 TFs represented in the sigma factor network. Therefore, our four TFs of interest were significantly enriched using hypergeometric test (p-value = 0.0027). According to the model, Lsr2 and DosR both interact with three sigma factors directly, whereas Rv0691c interacts with four sigma factors indirectly through their corresponding anti-sigma factors. To examine the potential of these three TFs of regulating the whole sigma factor network, we inferred the hierarchical organization of the sigma factor network using the hierarchy score maximization algorithm ^67^. This algorithm shows how a given directed network is organized in multiple levels and elucidates the position of each component at the inferred levels. The results indicated that the sigma factor network has either three or four levels, and that DosR, Lsr2 and Rv0691c fall into the top-level of the network propagating regulations towards the lower levels (Table S2). Moreover, SigB, controls the transcription factor DosR expression ^66^. Of note, only the protein RsfB (anti-anti-sigma factor) showed significant differential expression between the two lineages among all the sigma factors and their corresponding anti-sigma factors and anti-anti sigma factors quantified on transcript and protein level (Table S1). Overall, these three TFs located at the top-level of the sigma factor network interact with eight sigma factors that could, in part, facilitate stress responses of L2 strains compared to L1. As the DosR regulon showed a higher basal expression in L2 strains, we next considered the extent to which DosR proteins would respond differently to a relevant stress.

### Differential proteomic and phenotypic response of L1 and L2 strains to nitric oxide stress

To determine whether the molecular differences observed between L1 and L2 strains, such as in the DosR regulon, are functionally relevant, we exposed the strains to nitric oxide (NO) stress for 24 hours and analyzed the samples with respect to their growth status and protein expression levels. NO stress simulates the physiological environment that the bacilli experience following engulfment by macrophages ^68^. Specifically, we (i) quantified DosR target proteins and (ii) verified the growth arrest duration after NO stress (see Materials and Methods). The expression of DosR target proteins increased more strongly in response to NO in L2 strains compared to L1 (Wilcoxon signed rank test’s p-value = 0.00022) (Fig 6a). Given the tight interaction with DosR and sigma factors, we also examined differential expression of sigma factors in L1 and L2 on treatment with NO. Our differential protein expression analysis showed that the strength of SigB induction in response to NO differed between L1 (fold change = 1.24, adjusted p-value = 0.006) and L2 (fold change = 1.94, adjusted p-value = 3.37E-6) strains, while other quantified sigma factors remained unchanged in both lineages (Table S3). This observation highlights a potential molecular mechanism underlying the stronger induction of DosR proteins in L2 strains following NO stress compared to L1 strains. Altogether, DosR proteins showed not only a higher basal expression but also a stronger induction following NO challenge in L2 strains when compared to L1.

**Figure 6.**
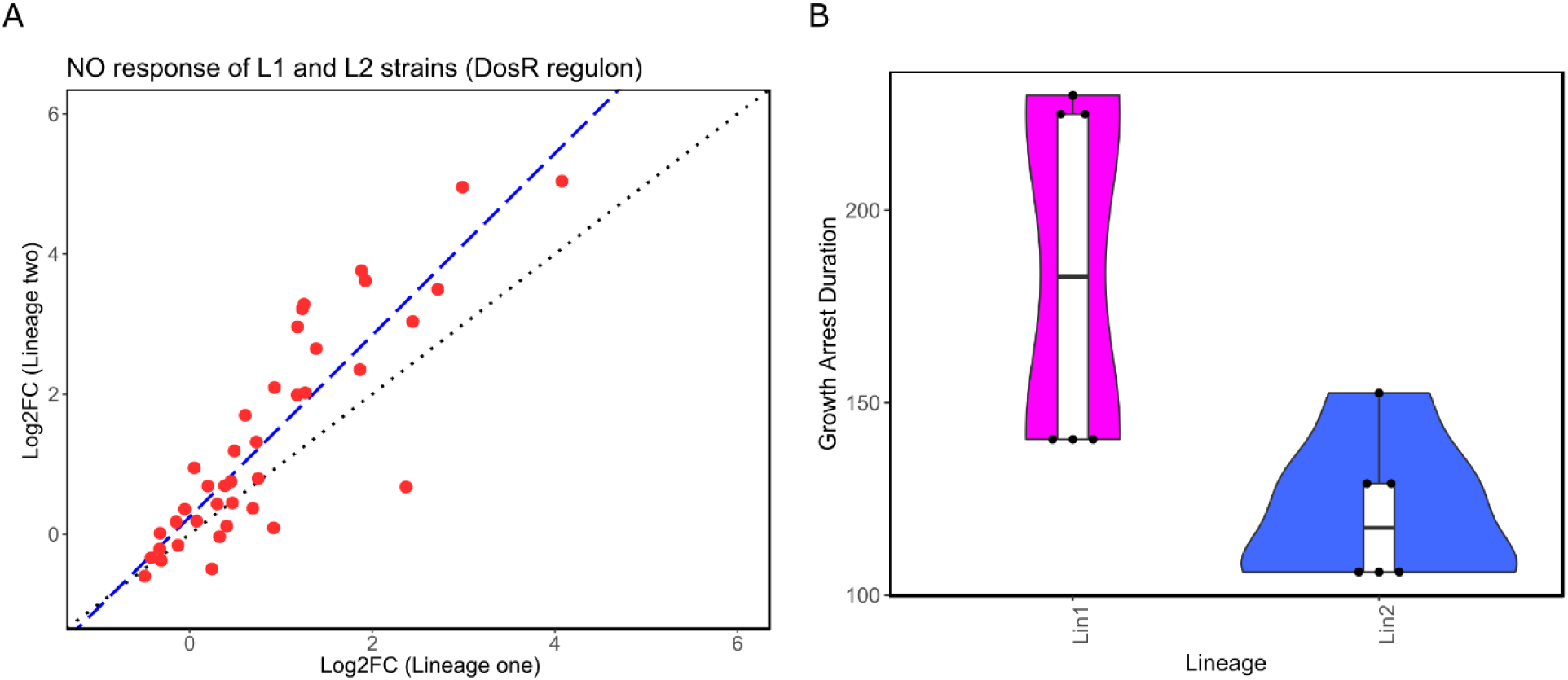
DosR regulon response to nitric oxide exposure and its phenotypic consequence. **(A)** Scatter plot shows bio-molecular responses of DosR proteins in MTBC clinical strains. Each dot represents a DosR protein while its respective response in L1 and L2 strains are denoted on X and Y axis respectively suggesting L2 strains responded significantly stronger to nitric oxide compared to L1 (Wilcoxon signed rank test’s p-value = 0.00022). **(B)** Shorter growth arrest duration upon NO exposure in L2 strains compared to L1 upon nitric oxide exposure (Wilcoxon signed rank test’s p-value = 0.0181).

Based on these molecular data, we hypothesized that following NO exposure, L2 strains, would be better capable to restart their growth compared to L1 strains. We tested this hypothesis by measuring the duration of growth arrest in two strains from each lineage in biological triplicates. The results demonstrated a significantly shorter growth arrest in L2 strains compared to L1 (Wilcoxon signed rank test’s p-value = 0.0181), supporting our hypothesis (Fig 6b). In summary, our findings show that TF network-based analysis can generate mechanistic hypotheses that are not directly apparent from the multi-omics data. We then validated the network-driven hypothesis experimentally and found that differential regulation of bio-molecular networks might drive distinct phenotypes in MTBC clinical isolates.

### Rv1985c induction accounts for one-third of the expression differences between L1 and L2 strains

Among the four master TFs identified by our transcriptional model, Rv1985c has been largely understudied in the scientific literature, and we could not leverage publicly available data to highlight its biological significance. However, we consider it a master regulator differentiating the two lineages for three reasons: first, it was identified by GenSTrans; second, its affected subnetwork, comprising 58 genes, was the largest (Fig 5a); and third, it is heavily involved in regulating DosR regulon genes (Fig 5e). Hence, we conducted a transcriptional regulator-induced phenotype (TRIP) ^69^ screen in the H37Rv background, in which ∼200 Mtb transcription factors were individually induced, to examine the extent to which each of Mtb TFs reveals phenotypes under hypoxia and normoxia. The results indicated that only a handful (∼5 TFs) of TFs, including Rv1985c, showed a significant phenotype (altered growth rate) when the two conditions were compared. These screening results go beyond the scope of this study. Nevertheless, we designed and implemented a time-course experiment in which the Rv1985c induction and control (empty plasmid) strains were induced one day prior to the experiment, followed by 7 days of hypoxia and 5 days of reaeration (Fig 7a). We measured CFU on days 0, 7, 8, 9, 10, 11, and 12, and performed RNASeq analysis on days 0, 2, 4, 7, 8, 9, 10, 11, and 12 (Fig. 7a, S9). The CFU results revealed a significant difference in the growth rate of the strains during the reaeration phase. Of note, Rv1985c is upregulated under hypoxia, and therefore this difference in growth rate was not expected during the hypoxic phase (Fig. 7b). Next, we conducted differential expression analysis between the induced and control strains at each time point to examine how many genes were up- or downregulated as a result of Rv1985c overexpression under different conditions. The results showed that approximately 500 genes were upregulated and 500 downregulated (fold change > 1.5 and adjusted p-value < 0.01) during hypoxia, and around 1,000 genes were upregulated and 1,000 downregulated during the reaeration phase (Fig. 7c; Table S4). Given that Rv1985c displayed significantly higher basal expression in the control strain under hypoxia compared to reaeration, the more modest transcriptional response to Rv1985c overexpression during hypoxia was expected.

**Figure 7.**
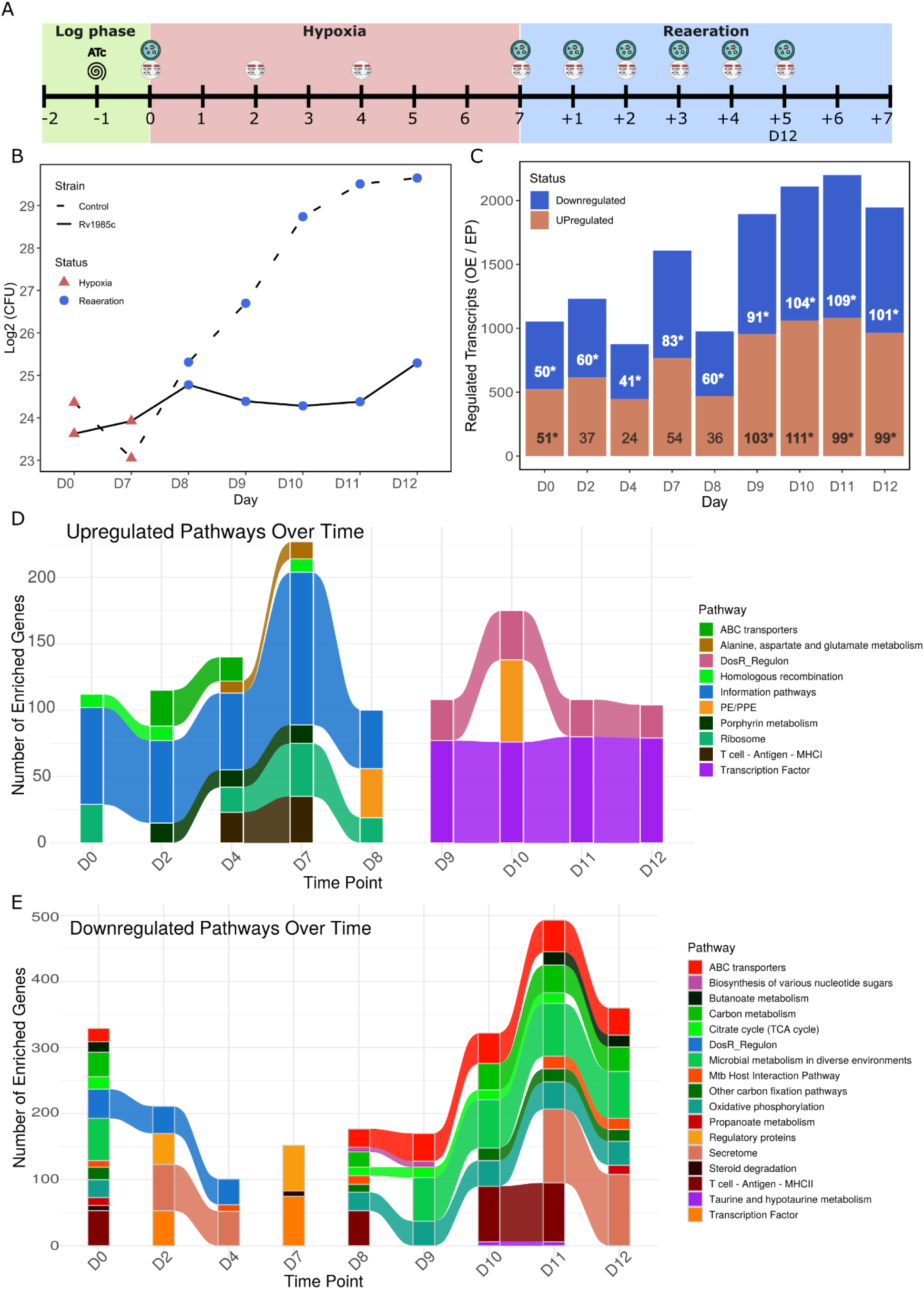
Rv1985c-mediated transcriptional and phenotypic responses to hypoxia and reaeration. **(A)** Schematic overview of the experimental design, indicating time points at which CFU counts and RNA-Seq measurements were performed. **(B)** Growth curves based on CFU counts of the Rv1985c induction and wild-type (empty plasmid) strains during hypoxia and reaeration phases. **(C)** Bar plots showing the number of significantly up- and downregulated genes in the induction strain compared to the wild type at each time point. Numbers on the bars indicate how many L2-vs-L1 upregulated (out of 275) and downregulated (out of 303) genes overlap with Rv1985c-regulated genes. Asterisks denote statistically significant enrichments based on the hypergeometric test. **(D, E)** Alluvial plots representing significantly enriched pathways among Rv1985c-upregulated **(D)** and -downregulated **(E)** genes across time points.

We next asked how many of the genes regulated by Rv1985c overlap with those differentially expressed between L2 and L1. For example, 51, 37, 24, 54, 36, 103, 111, 99, and 99 L2-upregulated genes (out of 275) were also upregulated on D0, D2, D4, D7, D8, D9, D10, D11, and D12, respectively (Fig. 7c). Similarly, 50, 60, 41, 83, 60, 91, 104, 109, and 101 L2-downregulated genes (out of 303) were also downregulated at the same time points (Fig. 7c & Fig. S10). We further performed hypergeometric testing to evaluate whether the L2/L1 up- and downregulated gene sets were significantly enriched within the Rv1985c-regulated sets at each time point. The results showed that the downregulated L2/L1 genes were significantly enriched at all time points, while the upregulated genes were enriched in nearly all time points during the normoxic/reaeration phase, with the exception of D8 (Table S4). Overall, these results validated that Rv1985c is indeed a master regulator differentiating L2 from L1 strains, explaining approximately one-third of the significant transcriptomic changes (∼200 out of 578 genes)—a substantially higher proportion than what was inferred from the transcriptional model alone. A possible explanation for this higher proportion could be that DosR and Rv1985c appear to share some targets, making it difficult to disentangle the individual contribution of each TF, especially since Rv1985c also affects DosR.

We further explored the function of the Rv1985c up- and downregulated genes at each time point to identify which Mtb pathways were enriched. We retained only significantly enriched pathways (adjusted p-value < 0.05) that appeared in at least two time points. Major upregulated pathways enriched in the Rv1985c-upregulated gene sets during hypoxia included “Information pathways” and “ABC transporters,” while “DosR regulon” and “Transcription factor” categories were highly enriched during the reaeration phase. This could explain why Rv1985c overexpression significantly altered ∼2,000 genes during reaeration. It appears that Rv1985c is an upstream TF that upregulates ∼80 other TFs. These findings further support the notation that Rv1985c and DosR work closely together, in line with previously published ChIP seq data ^61^. (Fig 5e, Fig 7d). Interestingly, the “DosR Regulon” was also significantly enriched among Rv1985c-downregulated genes during hypoxia. In addition, several metabolic pathways, such as the “TCA cycle” and “Microbial metabolism in diverse environments,” were significantly downregulated during reaeration (Fig 7e). This observation is further supported by a recent study that demonstrated reduced intracellular ATP levels in L2 strains compared to L1 strains ^70^. This reduced metabolic activity may contribute to increased drug tolerance, which could help explain why L2 strains are more associated with drug tolerance - such as to bedaquiline - compared to L1 strains ^70^. Overall, this time-course experiment validated Rv1985c as a master regulator differentiating the two lineages, demonstrated its close functional association with DosR, and provided a potential explanation for the increased drug tolerance observed in L2 strains.

## Discussion

We used a multi-omics approach combined with network analyses and validation experiments to compare clinical strains of the MTBC belonging to L1 and L2 to better understand what causes the differences in clinically relevant phenotypes such as virulence, transmission and drug resistance that have been observed between these lineages (Borrell et al., 2019). For this, we went beyond the conventional multi-layer omics data analysis approaches and introduced a new integrative, network-based analysis. Our results revealed that (i) the genetic distance between MTBC strains, based on SNPs, largely determines the number of significant quantitative changes at both the mRNA and protein levels; (ii) the extent of post-translational regulation in the MTBC varies across functional categories; (iii) compared to L1 strains, L2 strains exhibit reduced post-translational regulatory potential for genes annotated with “regulatory” and “virulence” functions; (iv) four transcription factors account for 25% of the expression differences observed between the two lineages; (v) these molecular changes translate into relevant phenotypes, such as reduced nitric oxide stress-induced growth arrest in L2 strains; and (vi) Rv1985c appears to play a major regulatory role distinguishing L1 from L2 strains, potentially explaining the increased bedaquiline tolerance observed in L2 ^70^.

First, we sought to elucidate to what extent SNPs cause significant changes at the mRNA and protein level. Hence, we correlated genomic distance (i.e. the number of distinct SNPs between a given pair of strains) and the number of significant changes on both transcript and protein levels. We showed that every third SNP on average is responsible for one significant change in gene expression. This observation could trigger a large community effort to map genotype – proteotype association in Mtb via protein quantitative trait locus (pQTL) analysis. This type of study has been implemented in various organisms ranging from yeast to mouse to human but not yet in bacterial systems ^71–73^.

Second, we investigated the significance of post-translational regulation, both systems-wide and for different functional categories. The analyses revealed that the degree of MTBC post-translational regulation as determined by the correlation between transcript and protein measurements resembles the patterns observed in higher organisms ^43^. While for some functional categories, such as lipid metabolism, the transcript – protein correlation was relatively high, for others such as virulence genes, it was very low. Therefore, transcriptomic measurements in isolation might be misleading in MTBC. For instance, Mtb regulates thousands of its transcripts in response to nitric oxide but only a few hundred of these changes are detectable on the protein level and the rest of them are buffered out ^20^. Our analyses revealed that for virulence and regulatory genes, post-translational events were less pronounced in L2 strains when compared to L1.

Third, we examined the extent to which significant changes between L1 and L2 strains can be explained by transcription factors (TFs). We started with the published EGRIN model, but this pipeline was insufficient to identify all causative TFs for two reasons, namely, the lack of specificity and its sparse coverage of TFs. Hence, we devised a genome-scale transcriptional network analysis, GenSTrans, which recapitulated the vast majority of TFs identified by the EGRIN model. It also shed some new light on 17 further TFs that were not represented in the EGRIN model. The analysis resulted in four transcription factors – DosR, Rv1985c, Rv0691c and Lsr2 – being linked to a quarter of the significant observed changes in gene expression. These transcription factors were significantly enriched in the sigma factor network and can regulate seven sigma factors, reprogramming RNA polymerase to modulate its affinity to promoter sequences under various stresses and conditions. Hierarchical analysis of the sigma factor network showed that the master TFs identified in this study fell into the top-level organization of the sigma factor network. This creates numerous possibilities for L2 strains to respond differently to a given stress compared to L1. Of note, SigB was the only sigma factor known to control the expression of one of our proposed master TFs. Since our analyses highlighted the differential basal expression of DosR genes between L1 and L2 on both transcript and protein level, we hypothesized that DosR regulon proteins might reveal a similarly differential trend in response to a relevant stress. Hence, we tested and validated this hypothesis in NO stress replicating a physiological stress that the tubercle bacilli experience following uptake by macrophages. The involvement of the sigma factors was enriched in our transcriptional regulatory analysis, and, in particular, the previously shown SigB regulation of DosR, was supported by the finding that SigB displayed a stronger induction with respect to NO stress in L2 vs L1 strains. This is also consistent with previous findings showing that SigB knockout strains are deficient in a variety of stress responses ^74,75^. Overall, the data showed that DosR proteins hold not only a higher basal expression but also a stronger response to NO stress in L2 strains compared to L1. We further elucidated that this molecular behavior empowers L2 strains to restart their growth following exposure to NO more rapidly. Altogether, our network analysis identified four major TFs orchestrating a large fraction of differential gene expression between L1 and L2 strains that could drive distinct phenotypes.

Fourth, we designed and implemented a time-course experiment to confirm the role of Rv1985c as a master TF, which as to-date not been thoroughly characterized. Using an inducted strain, we found that (i) Rv1985c is one of the few transcription factors showing a relevant phenotypic effect, (ii) Rv1985c plays a major role in differentiating L1 and L2 strains, explaining up to a third of the transcript-level changes, (iii) Rv1985c functions as an upstream regulator and shares several targets with DosR, and (iv) Rv1985c reduces metabolic activity during the reaeration phase, potentially explaining the molecular basis for the increased bedaquiline tolerance observed in L2 strains ^70^.

In summary, this study further highlights the fact that despite of its reduced genomic diversity compared to other bacteria, the MTBC phenotypically diverse. We showed that in clinical isolates of MTBC L1 and L2, hundreds of mRNAs and proteins are differentially expressed and that these molecular differences are being translated to phenotypes likely to be of relevance in the clinic. This study also provided important insights to guide future efforts to link specific genetic loci with mRNA and/or protein-level changes. Such a map of genetic features and their effect on molecular and macroscopic phenotypes in the MTBC provides an alternative regulatory archetype to those published; one derived from a genetically fairly conserved bacterium which may serve as a valuable counter point for future studies.

## Materials and Methods

### Mtb strains and bacterial culture

We used six clinical isolates of Mtb, three from L1 and three from L2, that are part of a recently described set of reference strains ^40^. These strains were chosen in an attempt to capture the global genetic diversity of Mtb, both within and between lineages. We grew bacteria in a modified 7H9 medium supplemented with 0.5% w/v pyruvate, 0.05% v/v tyloxapol, 0.2% w/v glucose, 0.5% bovine serum albumin (Fraction V, Roche) and 14.5 mM NaCl. With respect to conventional 7H9 culture medium, we omitted glycerol, tween 80, oleic acid and catalase. For global expression profiling experiments we cultured the bacteria in 1l bottles containing large glass beads to avoid clumping and 100 ml of media. We incubated the cultures at 37°C and rotated them continuously on a roller. For nitric oxide stress experiments we grew the bacteria in 50ml conical screwcap tubes containing either 17ml (proteomic profiling) or 10ml of culture medium. The cultures were incubated at 37°C on an orbital shaker.

### Transcriptomic profiling

We transferred a 40 ml aliquot of bacterial culture in mid-log phase (OD600 = 0.5 ± 0.1) into a 50ml Falcon conical tube containing 10 ml ice. We harvested the cells by centrifugation (3,000×g, 7 min, 4°C), re-suspended the pellet in 1 ml of RNApro solution (MP Biomedicals) and transferred the suspension to a Lysing matrix B tube (MP Biomedicals). We disrupted the bacterial cells using a FastPrep24 homogeniser (40s, intensity setting 6.0, MP Biomedicals). We clarified the lysate by centrifugation (12,000×g, 5 min, 4°C), transferred the supernatant to a clean tube and added chloroform. We separated the phases by centrifugation (12,000×g, 5 min, 4°C) and precipitated the nucleic acids from the aqueous phase by adding ethanol and incubating at −20C overnight. We performed a second acid phenol extraction to enrich for RNA. We treated our samples with DNAse I Turbo (Ambion), and removed stable RNAs by using the RiboZero Gram Positive ribosomal RNA depletion kit (Epicentre). We prepared the sequencing libraries using the TruSeq stranded Total RNA kit (Illumina) and sequenced them on a HiSeq2500 run in high output mode (50 cycles, single end).

The resulting reads were mapped to the Mtb H37Rv reference genome using BWA (ver. 0.7.13); the resulting mapping files were processed with samtools (ver. 1.3.1). Per-feature read counts were performed using the Python module htseq-count (ver. 0.6.1p1) and Python (ver. 2.7.11). We performed differential expression analysis using the R package edgeR and R (ver. 3.4.0) to identify Lineage-specific transcriptional changes.

### Environment and gene regulatory influence network (EGRIN) based transcriptional model

We used the EGRIN model to analyze our transcriptional data in the exact same way that described before ^59^. The model describes a set of modules that each one contains a dozen of co-regulated genes. The ChIPSeq and TFOE data paved the way to correspond each module to its potential regulators namely transcription factors (TFs) ^58^. Here, we projected our significant differentially regulated mRNAs (fold change > 1.5 and adjusted p-value < 0.01) between L1 and L2 strains onto the modules to figure out how significantly each module was enriched. The enrichment analysis was performed using hypergeometric test followed by Benjamini-Hochberg (BH) multiple testing correction (adjusted p-value < 0.05). To visualize a given identified TFs, we included all its corresponding modules even if they were not significantly enriched.

### Genome-scale transcriptional model and networks integration

To reconstruct the Mtb transcriptional network, we retrieved a ChIPSeq data publicly available at the Mtb portal V2 ^61^. We included both operon and direct interactions in the final network, which offered a great framework for genome-scale analysis. In contrast to the EGRIN model, the TFOE data was totally excluded as they largely feature indirect effects of a transcription factor overexpression to the Mtb transcriptome. The reconstructed network comprised 143 transcription factors and 2,943 genes and can easily be extended upon the availability of ChIPSeq data for the remaining transcription factors. Next, a new set of regulons/modules, each contained the target genes of a given TF, were defined. Such a definition provided a one-to-one relationship between the modules and the TFs. Moreover, the average size of each module became ∼65 genes, five times that of the EGRIN model, offering a larger specificity. The significantly regulated transcripts in L2 strains compared to L1 were tested against each module using a hypergeometric test followed by Benjamini-Hochberg (BH) multiple testing correction (adjusted p-value < 0.05). The identified TFs were compared to the EGRIN model results (see the color codes in the corresponding figures). Four TFs with the largest regulated gene size explaining 25% of the significant changes were introduced as the master regulators. This approach is easily portable to other bacterial systems upon the availability of respective ChIPSeq data. The TFs of the identified sub-network was integrated into the Mtb sigma factors network constructed by Gennaro and colleagues ^66^. This networks integration revealed the interactions between our four identified TFs (three of these were presented in the sigma factor network) and the Mtb sigma factors. Sigma factors can reprogram RNA polymerases and change their affinity to a given promoter. Hence, such a network integration showed how significantly the identified TFs are capable to present different phenotypic features in response to a perturbation. Notably, SigB as the only sigma factor which could modulate the transcription factor DosR provided a great opportunity to validate the capability such analyses.

### Proteomic profiling using DIA-MS

We aliquoted 20 OD equivalents from mid-log phase (OD_600_ = 0.5 ± 0.1) bacterial cultures (e.g. 40ml of OD_600_ 0.5) into 50ml conical tubes. In the case of nitric oxide stress we harvested 8ml of bacterial culture. We harvested the bacteria by centrifugation (3,000×g, 7 min, 4°C) and washed the pellet twice with cold PBS to remove tyloxapol from the samples. The washed pellets were re-suspended in lysis buffer, which contained 0.1M ammonium bicarbonate, 8M urea and 0.1% RapiGest (#186001861, Waters). We then transferred the suspension to a Lysing matrix B tube (MP Biomedicals) and disrupted the bacterial cells using a FastPrep24 homogeniser (40s, intensity setting 6.0, MP Biomedicals). We clarified the lysate by centrifugation (12,000×g, 5 min, 4°C) and sterilized it by filtering it twice through a 0.22 μm syringe filter (Milipore) prior to further processing.

Next, we measured the protein concentration of each sample using BCA assay (A53225, Thermo Fisher Scientific). Considering the measured concentrations, we started with 60 ug protein and appropriate volume of lysis buffer was added up to 100 ul to equalize the final concentration of each sample. Then, 5 mM tris(2-carboxyethyl)phosphine (TCEP) was added to reduce protein disulfide bonds while the sample were incubated at 37°C for 30 min. Afterwards, the free cysteine residues were alkylated by adding 40 mM iodoacetamide and incubating for 30 min in the dark. The samples were diluted 6 times with 0.05 M ammonium bicarbonate to reach a final Urea concentration below 2 M. To digest proteins, 1.2 ug sequencing grade modified trypsin (V5113, Promega) was added (w/w 1:50). The samples were incubated overnight (∼16 hours) at 37°C with gentle shaking of 300 rpm. To stop protein digestion, we acidified the samples (PH < 2) using formic acid followed by and incubation for 30 min with shaking of 500 rpm. We desalted the clear peptide solution using C18 MicroSpin columns (The Nest Group, 30-300 ug loading capacity). Next, we spiked in iRT peptides (Ki-3002, Biognosys) which allowed us to normalize retention time during the data processing step.

We measured the samples on a TripleTOF 5600 mass spectrometer (AB Sciex) coupled to a nanoLC system (Eksigent) as described before ^38^. The data acquisition was performed in DIA-MS mode using the 64 variable windows scheme.

The OpenSWATH workflow, containing three software tools, paved the way to process the data. We used the pan Mtb spectral library generated on the same type of mass spectrometry, TripleTOF 5600 ^37^. OpenSWATH extracted chromatograms and assigned peak groups according to the prior knowledge, the pan Mtb spectral library ^41^. Next, we performed PyProphet-cli to score the assigned peak groups using a semi-supervised algorithm. The data was filtered to global 1% FDR on both peptide and protein level. Eventually TRIC was used to integrate various information of each run and align the extracted and scored peak groups between runs offering a high degree of consistency between measurements ^76^. To perform absolute (based on top 3 peptides and top 5 transitions) and relative quantification, we used the R package SWATH2stats ^77^ and eventually aLFQ ^78^ and MSstats ^79^.

### Assesment of the response to nitric oxide stress

To measure the proteomic changes following nitric oxide (NO) exposure, we grew 17ml bacterial cultures to mid-log phase (OD_600_ approximately 0.5) and harvested 8ml of the culture for proteomic characterization. We then added Diethylenetriamine/NO adduct at a final concentration of 1mM to the remaining culture and incubated it for a further 24h at 37°C. After this time, we harvested 8ml of the culture for proteomic characterization.

To assess the duration of the NO induced growth arrest, we grew duplicate bacterial cultures to mid-log phase (OD_600_ approximately 0.5) and added Diethylenetriamine/NO adduct at a final concentration of 1mM to the treatment tube with nothing added to the control. We followed the subsequent growth dynamics by measuring OD_600_.

### Genetic, transcriptomic and proteomic distance analysis between clinical isolates

To compute strain distances on various levels, we assigned a particular cutoff to digitize them. On genome layer, we ignored various genetic insertions and deletions and only counted the numbers of distinct SNPs between a pair of Mtb strains. For mRNA and protein level, we considered a given cutoff (fold change > 1.5 and adjusted p-value < 0.01) and then the number of the significant changes were calculated. For each layer, 15 comparisons consisting of six intra lineage (three pairs within each lineage) and nine inter lineages comparisons were assumed. Afterwards Spearman correlation was computed between the genomic and either transcriptomic or proteomic distance. The slope of the respective regression line showed how many significant changes on protein and transcript level are caused by a given distinct SNP.

### Sigma factor network hierarchical properties

To infer the hierarchical organization of the sigma factor network recently described ^66^, we used the hierarchy score maximization algorithm ^67^. We performed the algorithm for different number of levels k (2-6). In each analysis, it computes probability scores to assign a given node to layers of the hierarchical organization. Two criteria were assessed to determine the optimal choice of k. First, the enrichment of the downward direction compared to expectation that is summarized in the corrected hierarchy score. Second, an ambiguity score which quantifies the uncertainty of a node assigning to a given layer of the organization. These two measurements revealed that k=4 is the optimal.

## Acknowledgements

We would like to thank Valerie F. A. March (Swiss TPH), John Aitchison (Seattle Children’s Hospital), Xueli Guan (Nanyang Technological University) and Uwe Sauer (ETH Zurich) for their intellectual contributions. We further thank Christoph Grundner (Seattle Children’s Hospital) for providing the PknH knockout and overexpressing strains. We would like to thank the Genomics Facility Basel for profiling the transcriptome of the clinical strains. We would also thank the Scientific IT Support (ID SIS) of ETH Zurich and the scientific computing center at University of Basel for support and maintenance of the laboratory-internal computing infrastructure. This work was supported by the SystemsX.ch project TbX, the National Institutes of Health project Omics4TB Disease Progression (U19 AI106761), the Swiss National Science Foundation (grants CRSII5_213514, 320030-227432 and 320030L-231163) and the European Research Council (883582-ECOEVODRTB). B.C.C. was supported by a Swiss National Science Foundation Ambizione grant (PZ00P3_161435).

## Author Contributions

Conceptualization: SG, ABE, BCC, RA, SB, and AT; Clinical strains cultivation: SB, SMG and JF; PknH strains cultivation: TRR and DRS; transcriptomic samples preparation: AT; Transcriptomic data acquisition: CB; Transcriptomic data processing and differential analysis: ABE, AT and SB; Proteomic samples preparation, data acquisition and data processing: ABE, BCC, LCG and OTS; Genome-scale transcriptional model: ABE; Growth arrest experiment: SB and JF; Other data analysis: ABE; Manuscript preparation: ABE (with critical inputs from SG, BCC and RA); Supervision: BCC, RA and SG.

## Data availability

The clinical strains and Rv1985c RNA-seq data are available on ArrayExpress under the dataset identifier E-MTAB-8103 and in the NCBI SRA under the BioProject accession PRJNA1297180. The mass spectrometry proteomics data have been deposited to the ProteomeXchange Consortium (PXD017194) via the PRIDE partner repository (Username, Password: zXJJJiHp).

---

**Figure S1.**
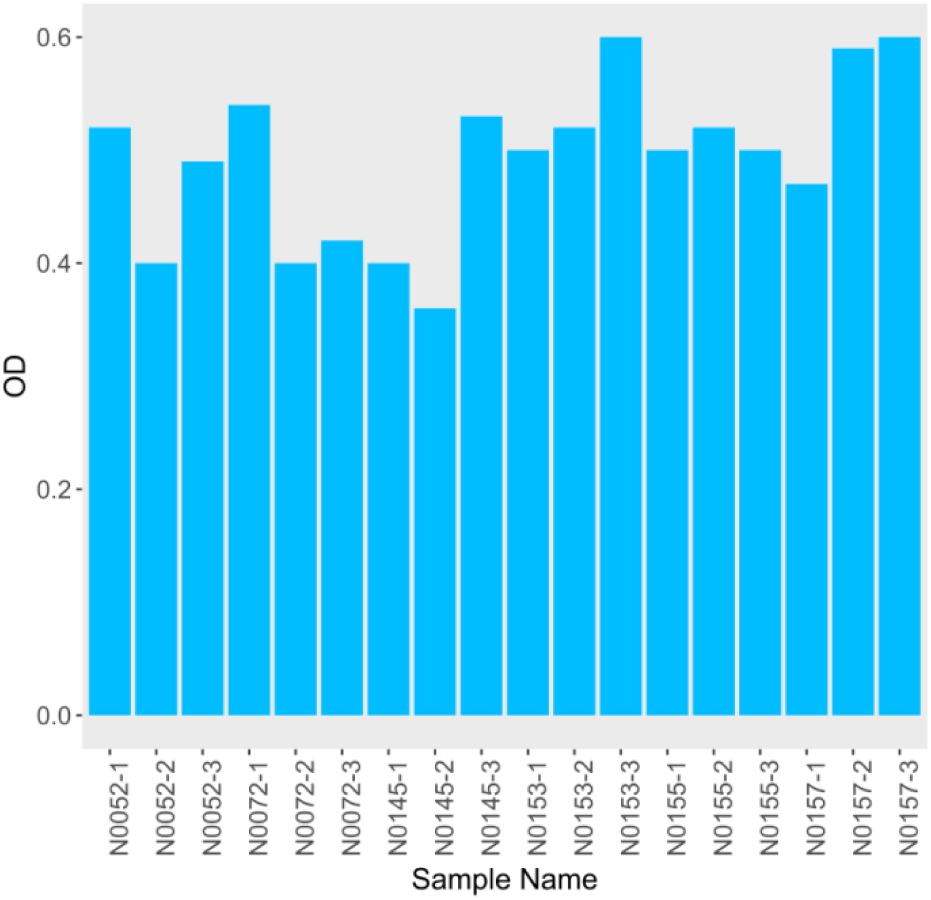
Bar plot of ODs measured at harvest time.

**Figure S2.**
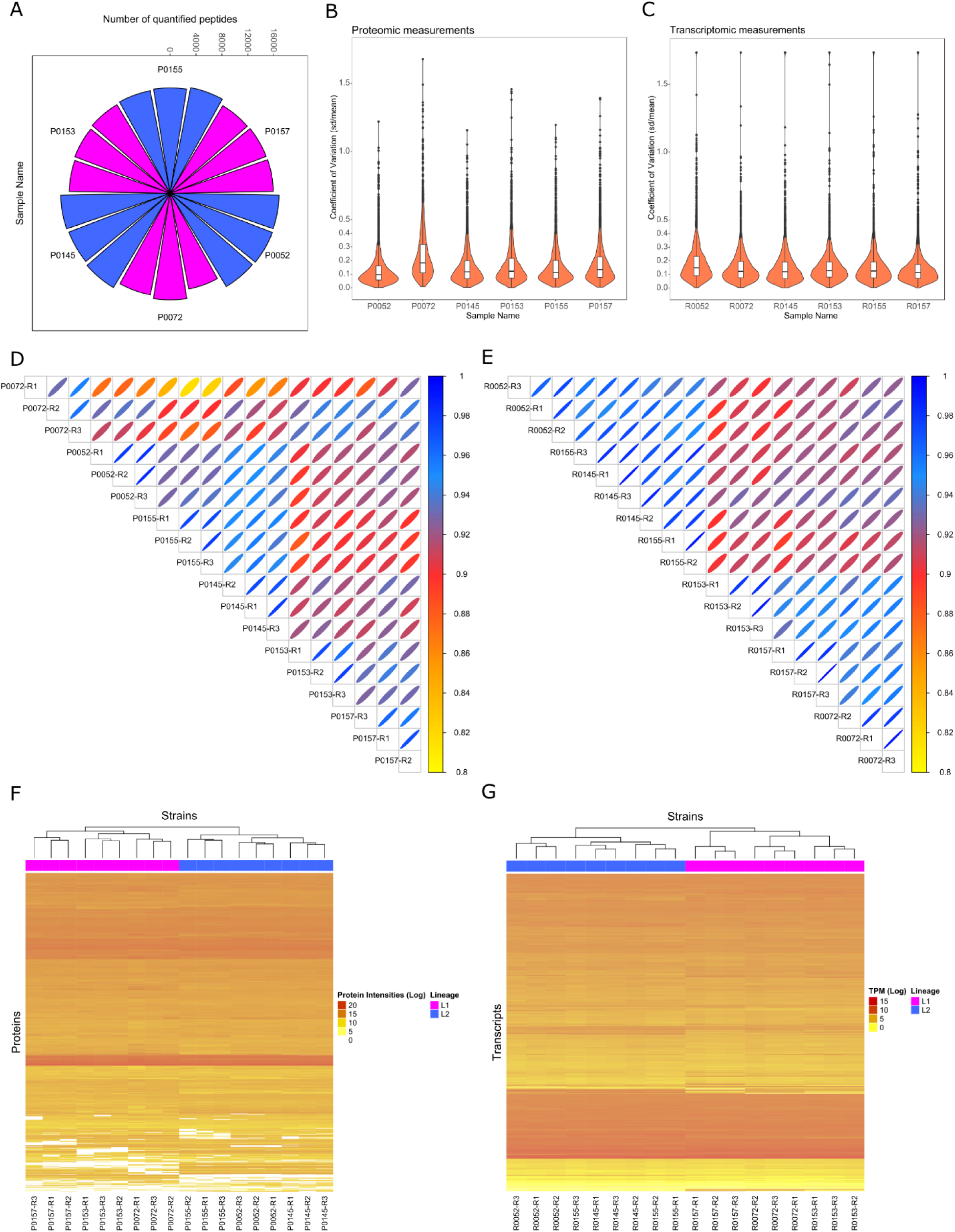
Transcript and protein data quality assessment. **(A)** Polar plot of peptide counts for each biological replicate of Mtb clinical strains. **(B-C)** Violin plots of CVs across biological replicates of each strains. The same median CVs (12%) suggested comparable reproducibility between Illumina NGS platform and DIA-MS used for transcriptomic and proteomic profiling of Mtb clinical strains. **(D-E)** Correlation plots of transcript (right) and protein (left) data. Spearman correlation analysis between clinical strains indicating near to perfect correlations and therefore reproducibility across biological replicates. **(F-G)** Perfect unsupervised clustering of clinical strains rooted in protein (left) and transcript (right) data.

**Figure S3.**
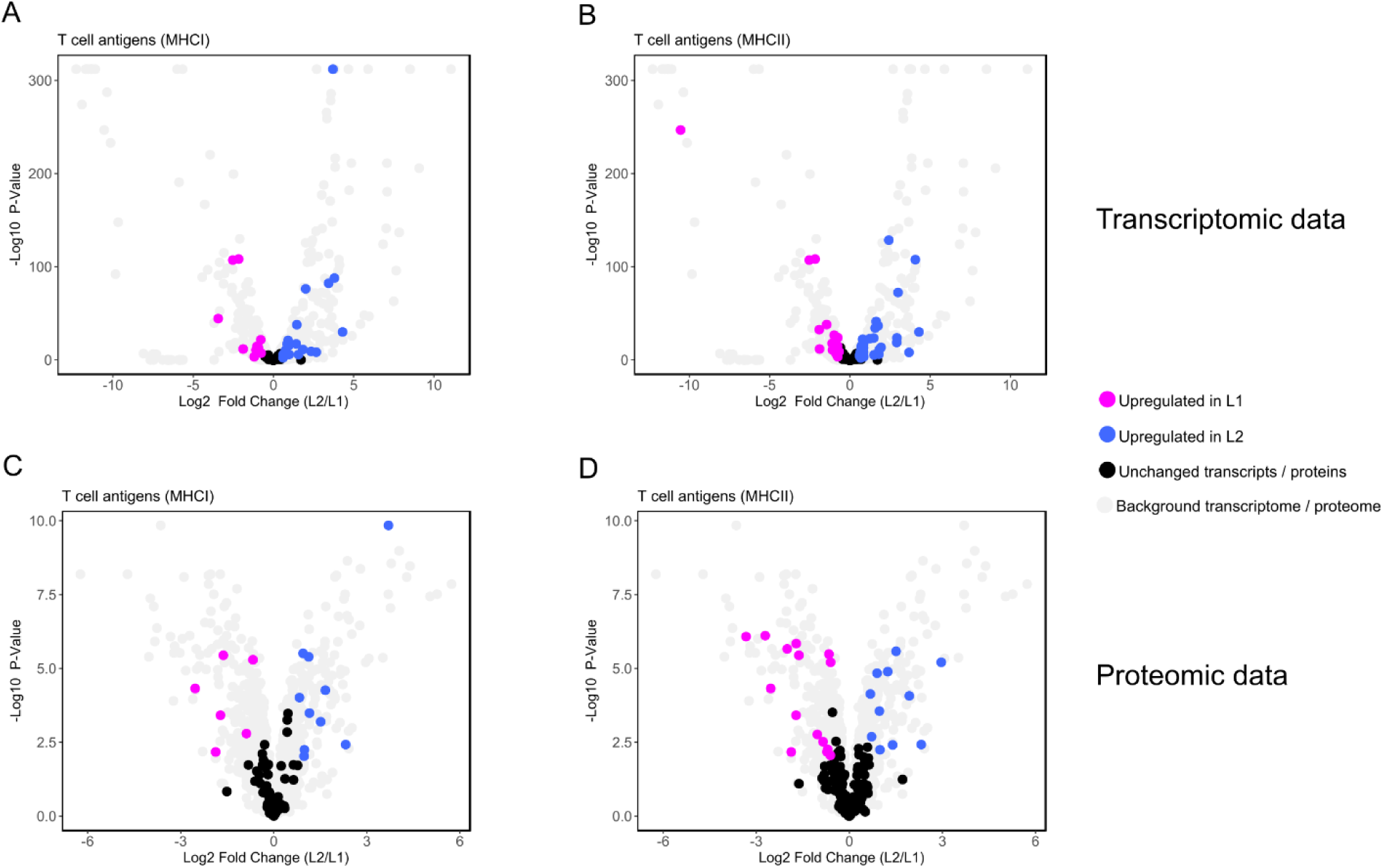
Enrichment analyses of T cell antigens. **(A-B)** Volcano plot of T cell antigens presented by MHCI (left) and MHCII (right) at the mRNA level. The analysis showed that these functional categories are significantly enriched (p-value for MHCI = 4.5E-5, p-value for MHCII = 0.0071) by the differentially expressed transcripts between the two lineages. **(C-D)** Enrichment analysis of differentially expressed proteins in L2 strains compared to L1. MHCI (left) and MHCII (right) presented antigens were not significantly enriched anymore on protein level (p-value for MHCI = 0.069, p-value for MHCII = 0.55).

**Figure S4.**
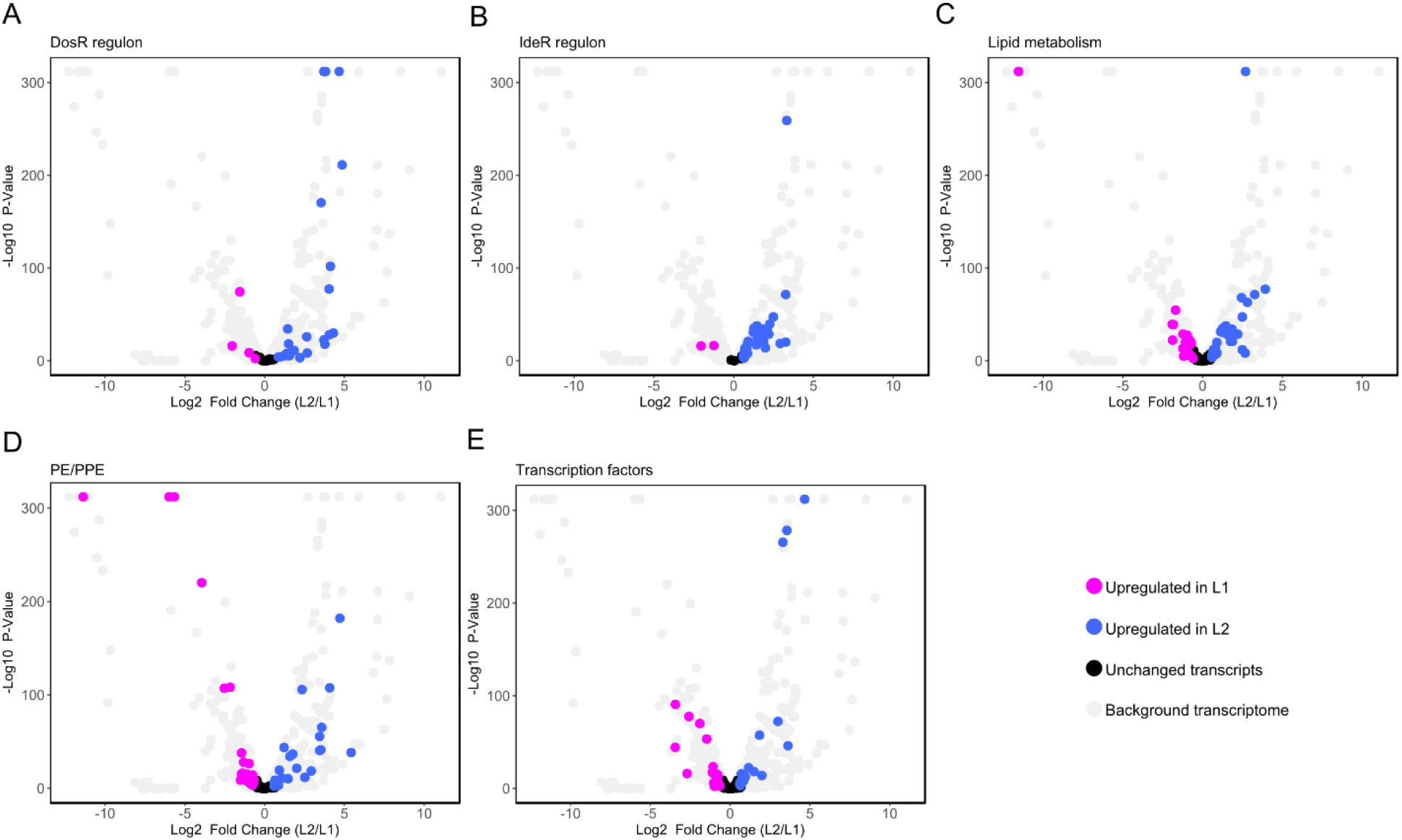
Enrichment analyses of MTBC functional categories by mRNA data. Volcano plots display that **(A)** DosR regulon (p-value = 4.08E-14), **(B)** IdeR regulon (p-value = 3.05E-28), **(C)** lipid metabolism (p-value = 0.0025), and **(D)** PE/PPE (p-value = 9.3E-9) are significantly regulated between L1 and L2 clinical strains at the RNA level. **(E)** Regulated transcription factors between the two lineages on mRNA level. 34 (out of 211 quantified) transcription factors were significantly regulated in L2 strains in respect to L1. (see Figure 3 for protein data)

**Figure S5.**
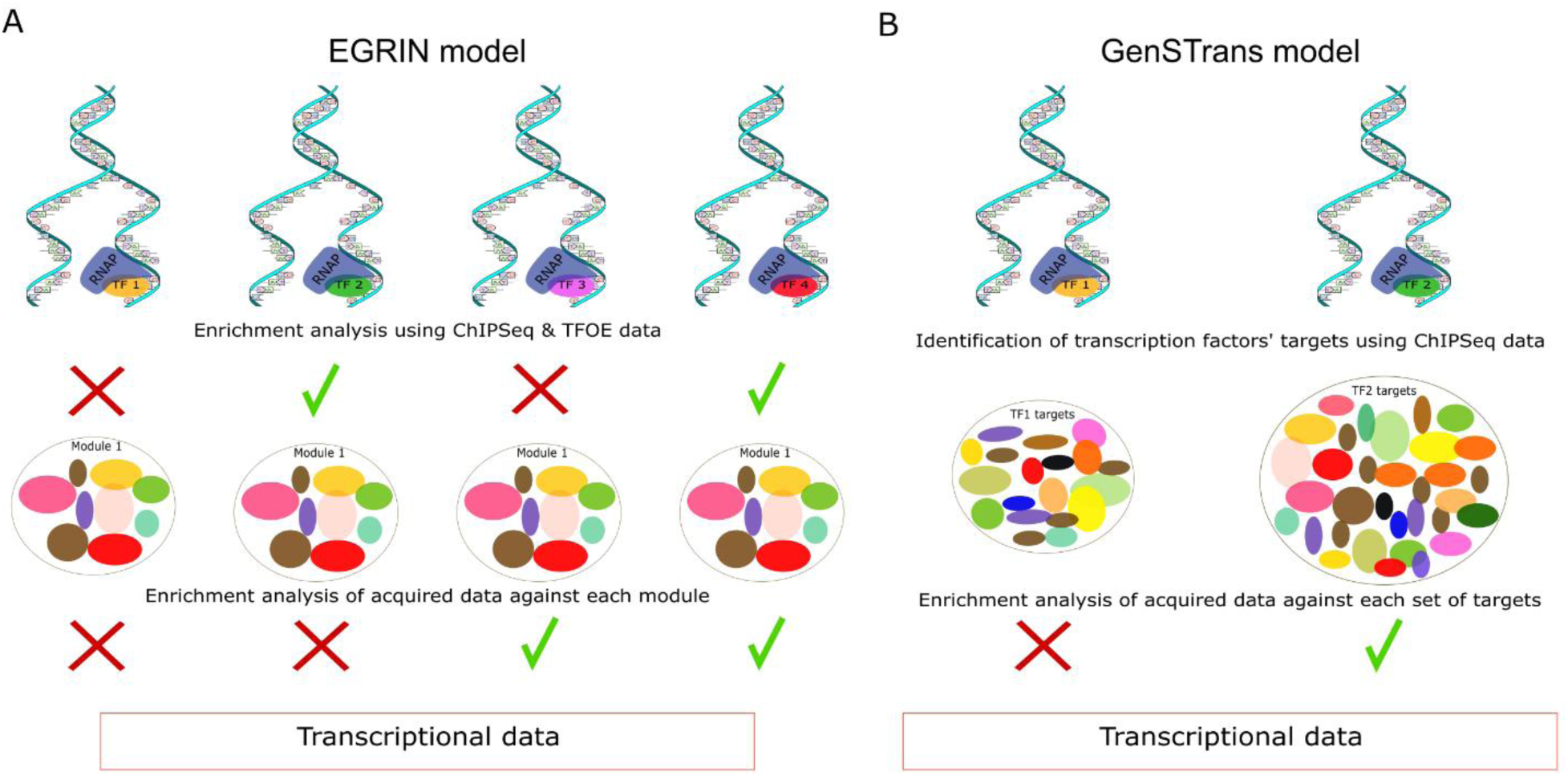
Conceptual diagrams for MTBC transcriptional modeling. **(A)** Schematic illustration of the EGRIN model. The EGRIN model is a module centric approach, which demonstrates clusters of co-regulated genes (module) inferred from hundreds of RNA profiles that are publicly available. Next, each module was linked to transcription factor(s) using both Transcription Factor OverExpression (TFOE) ^32^ and ChIPSeq ^61^ experiments by enrichment analysis. Therefore, some of modules could not be corresponded to any transcription factor and remained as orphans. Eventually acquired RNA data for a given study should be analyzed against modules. **(B)** Concept of GenSTrans. GenSTrans was developed using ChIPSeq data ^61^ and could be considered as a transcription factor centric model. This model explained sub-networks (instead of modules in the EGRIN model) which were reconstructed for each transcription factor. Therefore, each sub-network was linked to one and only one transcription factor. Finally, acquired RNA data could be analyzed through sub-networks.

**Figure S6.**
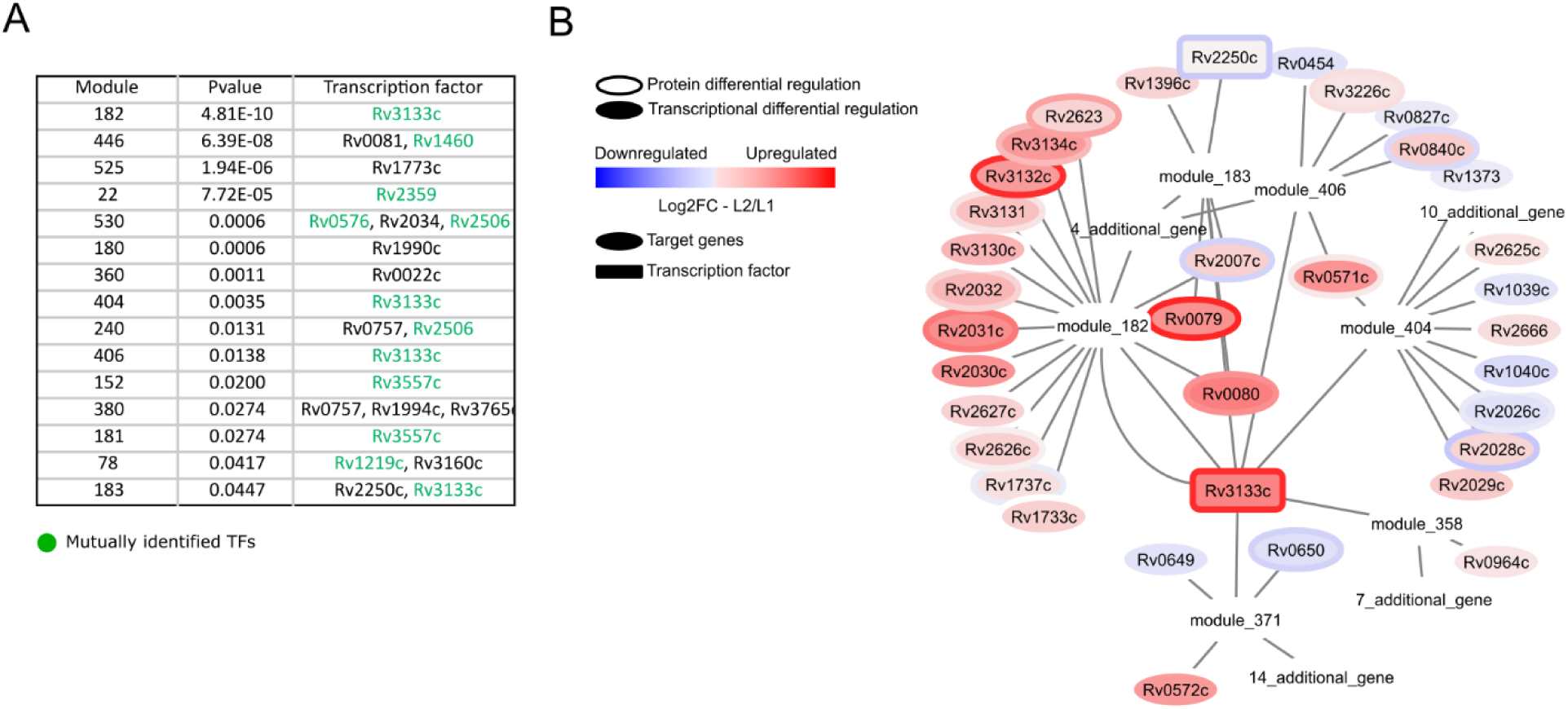
EGRIN based identification of transcription factors regulating their target modules differently between L1 and L2 Strains. **(A)** Identified modules and corresponding transcription factors by the EGRIN model. Green transcription factors were also identified by GenSTrans. **(B)** The transcription factor DosR that was identified through four modules significantly.

**Figure S7.**
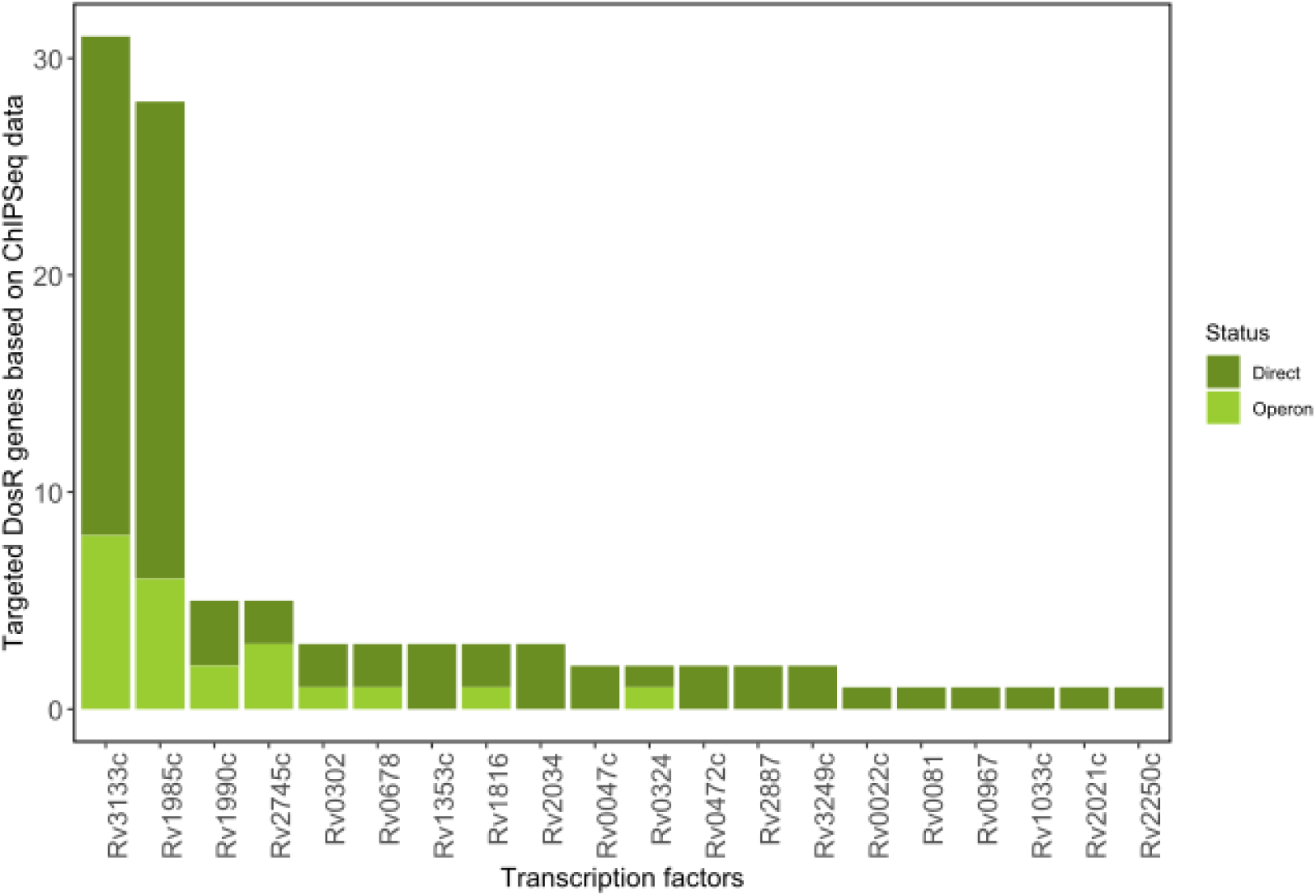
DosR regulon genes targeted by various transcription factors based on ChIP seq data.

**Figure S8.**
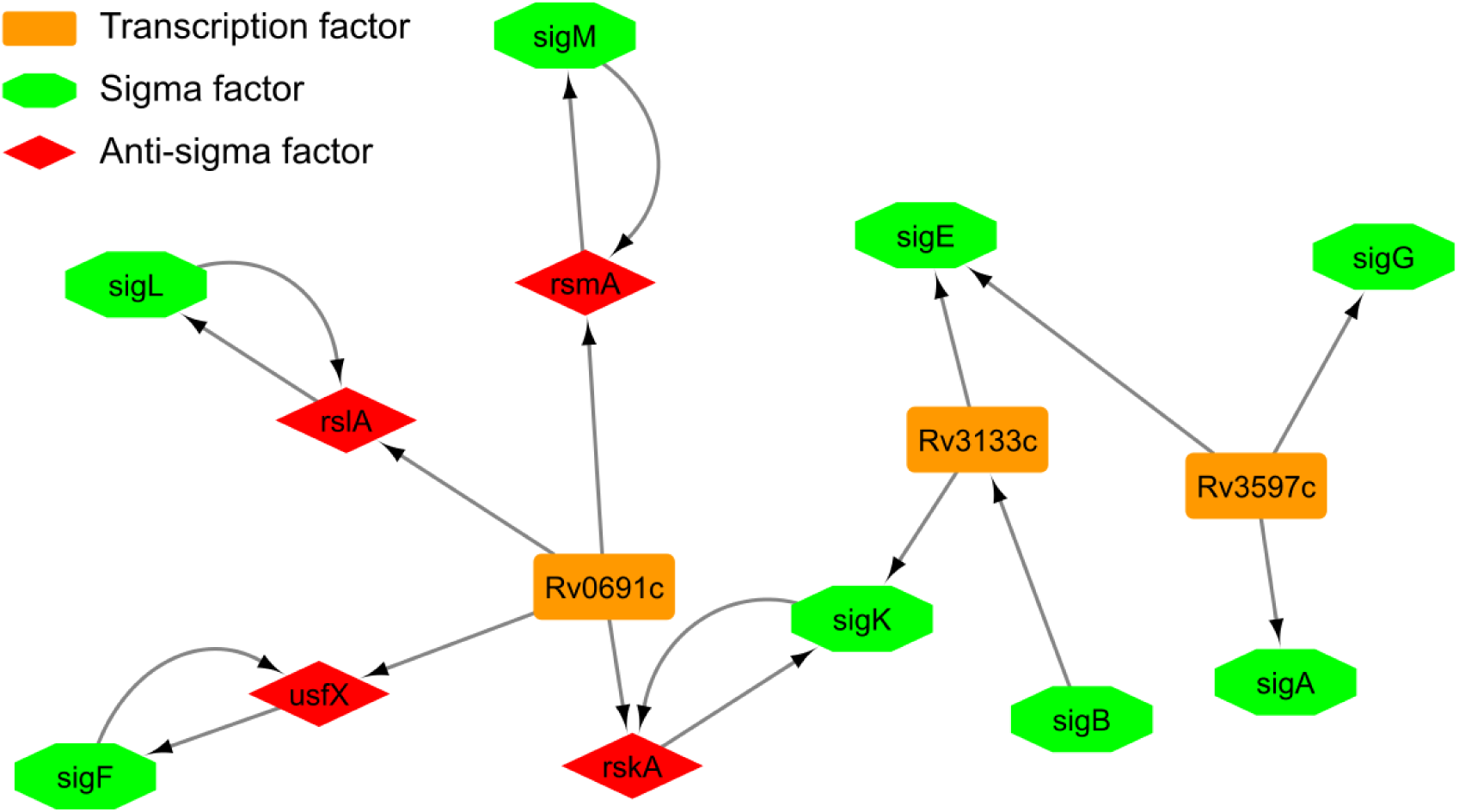
Interactions of inter-L1 and L2 master transcription factors with sigma factors. Network shows that each of Rv3133c (DosR) and Rv3597c interact with three sigma factors whereas Rv0691c could regulate four anti-sigma factor and therefore modulating sigma factors indirectly.

**Figure S9.**
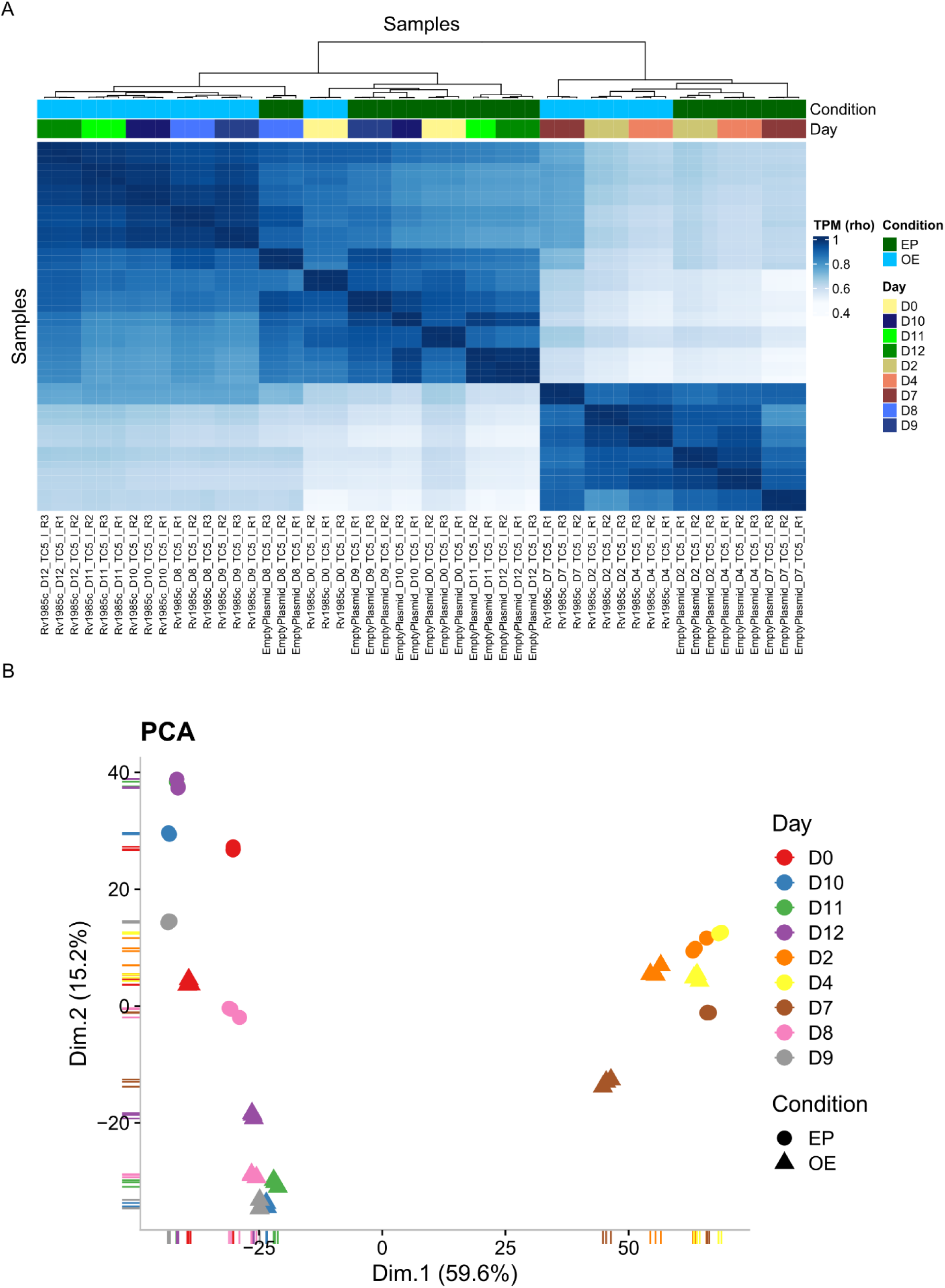
Overview of RNA seq data from the Rv1985c experiment. **(A)** Hierarchical clustering based on Spearman correlation of TPM values across all RNA-Seq runs. **(B)** Principal component analysis (PCA) plot of all RNA-Seq samples.

**Figure S10.**
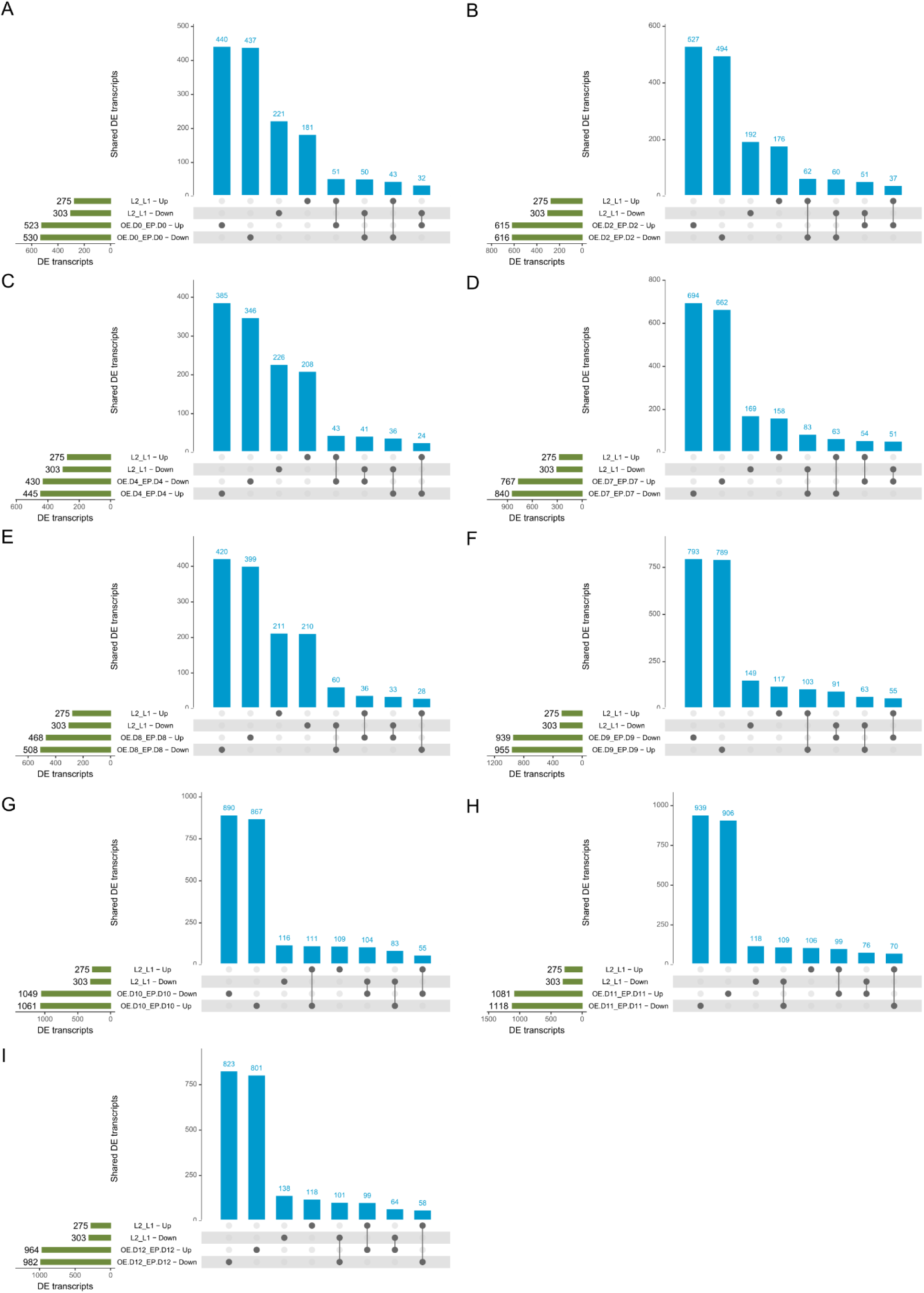
Comparison of Rv1985c-regulated genes with L2–L1 differentially expressed genes. Upset plots showing the overlap between Rv1985c up- and downregulated genes and L2–L1 up- and downregulated genes at each time point: **(A)** Day 0, **(B)** Day 2, **(C)** Day 4, **(D)** Day 7, **(E)** Day 8, **(F)** Day 9, **(G)** Day 10, **(H)** Day 11, and **(I)** Day 12.

## Reference

1. World Health Organization. Global Tuberculosis Report 2024 (WHO, 2024). https://www.who.int/teams/global-programme-on-tuberculosis-and-lung-health/tb-reports/global-tuberculosis-report-2024.

2. Goig, G. A. et al. Ecology, global diversity and evolutionary mechanisms in the Mycobacterium tuberculosis complex. Nature Reviews Microbiology 2025 1–13 (2025) doi:10.1038/s41579-025-01159-w.

3. Dartois, V. A. et al. Strategies for shortening tuberculosis therapy. Nat Med 31, 1765– 1775 (2025).

4. Achtman, M. Evolution, Population Structure, and Phylogeography of Genetically Monomorphic Bacterial Pathogens. Annu Rev Microbiol 62, 53–70 (2008).

5. Coscolla, M. & Gagneux, S. Consequences of genomic diversity in Mycobacterium tuberculosis. Semin Immunol 26, 431–44 (2014).

6. Arbués, A. et al. Soluble immune mediators orchestrate protective in vitro granulomatous responses across Mycobacterium tuberculosis complex lineages. Elife 13, (2025).

7. Brites, D. & Gagneux, S. The Nature and Evolution of Genomic Diversity in the Mycobacterium tuberculosis Complex. in Advances in experimental medicine and biology vol. 1019 1–26 (2017).

8. de Jong, B. C. et al. Progression to active tuberculosis, but not transmission, varies by Mycobacterium tuberculosis lineage in The Gambia. J Infect Dis 198, 1037–43 (2008).

9. Holt, K. E. et al. Frequent transmission of the Mycobacterium tuberculosis Beijing lineage and positive selection for the EsxW Beijing variant in Vietnam. Nat Genet 50, 849–856 (2018).

10. Borrell, S. & Gagneux, S. Infectiousness, reproductive fitness and evolution of drug-resistant Mycobacterium tuberculosis. Int J Tuberc Lung Dis 13, 1456–66 (2009).

11. Long, R. et al. The association between phylogenetic lineage and the subclinical phenotype of pulmonary tuberculosis: A retrospective 2-cohort study. Journal of Infection 88, 123–131 (2024).

12. Bhatia, A. L. et al. The virulence in the guinea-pig of tubercle bacilli isolated before treatment from South Indian patients with pulmonary tuberculosis. 2. Comparison with virulence of tubercle bacilli from British patients. Bull World Health Organ 25, 313–22 (1961).

13. Phyu, S. et al. Predominance of Mycobacterium tuberculosis EAI and Beijing Lineages in Yangon, Myanmar. J Clin Microbiol 47, 335–344 (2009).

14. Guerra-Assunção, J. et al. Large-scale whole genome sequencing of M. tuberculosis provides insights into transmission in a high prevalence area. Elife 4, (2015).

15. Coscolla, M. & Gagneux, S. Does M. tuberculosis genomic diversity explain disease diversity? Drug Discov Today Dis Mech 7, e43–e59 (2010).

16. Bottai, D. et al. TbD1 deletion as a driver of the evolutionary success of modern epidemic Mycobacterium tuberculosis lineages. Nat Commun 11, (2020).

17. Chitale, P. et al. A comprehensive update to the Mycobacterium tuberculosis H37Rv reference genome. Nature Communications 2022 13:1 13, 1–12 (2022).

18. Kalia, N. P. et al. M. tuberculosis relies on trace oxygen to maintain energy homeostasis and survive in hypoxic environments. Cell Rep 42, 112444 (2023).

19. Sawyer, E. B., Phelan, J. E., Clark, T. G. & Cortes, T. A snapshot of translation in Mycobacterium tuberculosis during exponential growth and nutrient starvation revealed by ribosome profiling. Cell Rep 34, 108695 (2021).

20. Cortes, T. et al. Delayed effects of transcriptional responses in Mycobacterium tuberculosis exposed to nitric oxide suggest other mechanisms involved in survival. Sci Rep 7, 8208 (2017).

21. Trauner, A. et al. Expression Dysregulation as a Mediator of Fitness Costs in Antibiotic Resistance. Antimicrob Agents Chemother 65, e0050421 (2021).

22. Wu, C. et al. Enzyme expression kinetics by Escherichia coli during transition from rich to minimal media depends on proteome reserves. Nat Microbiol 8, 347–359 (2023).

23. Mori, M. et al. From coarse to fine: the absolute Escherichia coli proteome under diverse growth conditions. Mol Syst Biol 17, e9536 (2021).

24. Caron, E., Aebersold, R., Banaei-Esfahani, A., Chong, C. & Bassani-Sternberg, M. A Case for a Human Immuno-Peptidome Project Consortium. Immunity 47, 203–208 (2017).

25. Nicod, C., Banaei-Esfahani, A. & Collins, B. C. Elucidation of host-pathogen protein-protein interactions to uncover mechanisms of host cell rewiring. Curr Opin Microbiol 39, 7–15 (2017).

26. Rose, G. et al. Mapping of genotype-phenotype diversity among clinical isolates of mycobacterium tuberculosis by sequence-based transcriptional profiling. Genome Biol Evol 5, 1849–62 (2013).

27. Comas, I. et al. Out-of-Africa migration and Neolithic coexpansion of Mycobacterium tuberculosis with modern humans. Nat Genet 45, 1176–82 (2013).

28. Heusel, M. et al. Complex-centric proteome profiling by SEC-SWATH-MS. Mol Syst Biol 15, e8438 (2019).

29. Bludau, I. et al. Complex-centric proteome profiling by SEC-SWATH-MS for the parallel detection of hundreds of protein complexes. Nat Protoc 15, 2341–2386 (2020).

30. Banaei-Esfahani, A., Nicod, C., Aebersold, R. & Collins, B. C. Systems proteomics approaches to study bacterial pathogens: application to Mycobacterium tuberculosis. Curr Opin Microbiol 39, 64–72 (2017).

31. Bei, C. et al. Genetically encoded transcriptional plasticity underlies stress adaptation in Mycobacterium tuberculosis. Nat Commun 15, (2024).

32. Rustad, T. R. et al. Mapping and manipulating the Mycobacterium tuberculosis transcriptome using a transcription factor overexpression-derived regulatory network. Genome Biol 15, 502 (2014).

33. Manson, A. L. et al. Genomic analysis of globally diverse Mycobacterium tuberculosis strains provides insights into the emergence and spread of multidrug resistance. Nat Genet 49, 395–402 (2017).

34. Buljan, M. et al. A computational framework for the inference of protein complex remodeling from whole-proteome measurements. Nat Methods (2023) doi:10.1038/s41592-023-02011-w.

35. Ludwig, C. et al. Data-independent acquisition-based SWATH-MS for quantitative proteomics: a tutorial. Mol Syst Biol 14, e8126 (2018).

36. Schubert, O. T. et al. Absolute Proteome Composition and Dynamics during Dormancy and Resuscitation of Mycobacterium tuberculosis. Cell Host Microbe 18, 96–108 (2015).

37. Schubert, O. T. et al. The Mtb Proteome Library: A Resource of Assays to Quantify the Complete Proteome of Mycobacterium tuberculosis. Cell Host Microbe 13, 602–612 (2013).

38. Collins, B. C. et al. Multi-laboratory assessment of reproducibility, qualitative and quantitative performance of SWATH-mass spectrometry. Nat Commun 8, 291 (2017).

39. Chiner-Oms, Á. et al. Genome-wide mutational biases fuel transcriptional diversity in the Mycobacterium tuberculosis complex. Nat Commun 10, (2019).

40. Borrell, S. et al. Reference set of Mycobacterium tuberculosis clinical strains: A tool for research and product development. PLoS One 14, e0214088 (2019).

41. Röst, H. L. et al. OpenSWATH enables automated, targeted analysis of data-independent acquisition MS data. Nat Biotechnol 32, 219–223 (2014).

42. Kapopoulou, A., Lew, J. M. & Cole, S. T. The MycoBrowser portal: A comprehensive and manually annotated resource for mycobacterial genomes. Tuberculosis 91, 8–13 (2011).

43. Liu, Y. et al. Multi-omic measurements of heterogeneity in HeLa cells across laboratories. Nat Biotechnol 37, 314–322 (2019).

44. Balakrishnan, R. et al. Principles of gene regulation quantitatively connect DNA to RNA and proteins in bacteria. Science (1979) 378, (2022).

45. Alli Shaik, A., et al. Functional Mapping of the Zebrafish Early Embryo Proteome and Transcriptome. J Proteome Res 13, 5536–5550 (2014).

46. Ghaemmaghami, S. et al. Global analysis of protein expression in yeast. Nature 425, 737– 741 (2003).

47. Brockmann, R., Beyer, A., Heinisch, J. J. & Wilhelm, T. Posttranscriptional Expression Regulation: What Determines Translation Rates? PLoS Comput Biol 3, e57 (2007).

48. Beyer, A., Hollunder, J., Nasheuer, H.-P. & Wilhelm, T. Post-transcriptional Expression Regulation in the Yeast *Saccharomyces cerevisiae* on a Genomic Scale. Molecular & Cellular Proteomics 3, 1083–1092 (2004).

49. Becker, S. H. et al. The Mycobacterium tuberculosis Pup-proteasome system regulates nitrate metabolism through an essential protein quality control pathway. Proc Natl Acad Sci U S A 116, 3202–3210 (2019).

50. Mehra, S. et al. The DosR Regulon Modulates Adaptive Immunity and Is Essential for *Mycobacterium tuberculosis* Persistence. Am J Respir Crit Care Med 191, 1185–1196 (2015).

51. Homolka, S., Niemann, S., Russell, D. G. & Rohde, K. H. Functional Genetic Diversity among Mycobacterium tuberculosis Complex Clinical Isolates: Delineation of Conserved Core and Lineage-Specific Transcriptomes during Intracellular Survival. PLoS Pathog 6, e1000988 (2010).

52. Reed, M. B., Gagneux, S., DeRiemer, K., Small, P. M. & Barry, C. E. The W-Beijing lineage of Mycobacterium tuberculosis overproduces triglycerides and has the DosR dormancy regulon constitutively upregulated. J Bacteriol 189, 2583–2589 (2007).

53. Chao, J. D. et al. Convergence of Ser/Thr and Two-component Signaling to Coordinate Expression of the Dormancy Regulon in *Mycobacterium tuberculosis*. Journal of Biological Chemistry 285, 29239–29246 (2010).

54. Domenech, P. et al. Unique regulation of the DosR regulon in the Beijing lineage of Mycobacterium tuberculosis. J Bacteriol 199, (2017).

55. Rodriguez, G. M. & Smith, I. Identification of an ABC Transporter Required for Iron Acquisition and Virulence in Mycobacterium tuberculosis. J Bacteriol 188, 424–430 (2006).

56. Rodriguez, G. M., Voskuil, M. I., Gold, B., Schoolnik, G. K. & Smith, I. ideR, An essential gene in mycobacterium tuberculosis: role of IdeR in iron-dependent gene expression, iron metabolism, and oxidative stress response. Infect Immun 70, 3371–81 (2002).

57. Gold, B., Rodriguez, G. M., Marras, S. A. E., Pentecost, M. & Smith, I. The Mycobacterium tuberculosis IdeR is a dual functional regulator that controls transcription of genes involved in iron acquisition, iron storage and survival in macrophages. Mol Microbiol 42, 851–865 (2008).

58. Peterson, E. J. R. R. et al. A high-resolution network model for global gene regulation in Mycobacterium tuberculosis. Nucleic Acids Res 42, 11291–303 (2014).

59. Peterson, E. J. R., Ma, S., Sherman, D. R. & Baliga, N. S. Network analysis identifies Rv0324 and Rv0880 as regulators of bedaquiline tolerance in Mycobacterium tuberculosis. Nat Microbiol 1, 16078 (2016).

60. Kendall, S. L. et al. Cholesterol utilization in mycobacteria is controlled by two TetR-type transcriptional regulators: kstR and kstR2. Microbiology (Reading) 156, 1362–71 (2010).

61. Minch, K. J. et al. The DNA-binding network of Mycobacterium tuberculosis. Nat Commun 6, 5829 (2015).

62. Frando, A. et al. The Mycobacterium tuberculosis protein O-phosphorylation landscape. Nature Microbiology 2023 8:3 8, 548–561 (2023).

63. Turapov, O. et al. Two Faces of CwlM, an Essential PknB Substrate, in Mycobacterium tuberculosis. Cell Rep 25, 57–67.e5 (2018).

64. Sachdeva, P., Misra, R., Tyagi, A. K. & Singh, Y. The sigma factors of Mycobacterium tuberculosis: regulation of the regulators. FEBS Journal 277, 605–626 (2010).

65. Rodrigue, S., Provvedi, R., Jacques, P.-É., Gaudreau, L. & Manganelli, R. The σ factors of *Mycobacterium tuberculosis*. FEMS Microbiol Rev 30, 926–941 (2006).

66. Chauhan, R. et al. Reconstruction and topological characterization of the sigma factor regulatory network of Mycobacterium tuberculosis. Nat Commun 7, 11062 (2016).

67. Cheng, C. et al. An approach for determining and measuring network hierarchy applied to comparing the phosphorylome and the regulome. Genome Biol 16, 63 (2015).

68. Nathan, C. Role of iNOS in human host defense. Science 312, 1874–5; author reply 1874-5 (2006).

69. Ma, S. et al. Transcriptional regulator-induced phenotype screen reveals drug potentiators in Mycobacterium tuberculosis. Nature Microbiology 2020 6:1 6, 44–50 (2020).

70. March, V. F. A. et al. Variability in intrinsic drug tolerance in Mycobacterium tuberculosis corresponds with phylogenetic lineage. *bioRxiv* 2025.07.04.663131 (2025) doi:10.1101/2025.07.04.663131.

71. Weith, M. et al. Genetic effects on molecular network states explain complex traits. Mol Syst Biol 19, (2023).

72. Li, H. et al. Integrative systems analysis identifies genetic and dietary modulators of bile acid homeostasis. Cell Metab 34, 1594–1610.e4 (2022).

73. Sobottka, B. et al. Immune phenotype-genotype associations in primary clear cell renal cell carcinoma and matched metastatic tissue. Mod Pathol 100558 (2024) doi:10.1016/j.modpat.2024.100558.

74. Manganelli, R. et al. σ Factors and Global Gene Regulation in Mycobacterium tuberculosis. Journal of Bacteriology vol. 186 895–902 Preprint at 10.1128/JB.186.4.895-902.2004 (2004).

75. Datta, P., Shi, L., Bibi, N., Balázsi, G. & Gennaro, M. L. Regulation of central metabolism genes of Mycobacterium tuberculosis by parallel feed-forward loops controlled by sigma factor E (σE). J Bacteriol 193, 1154–1160 (2011).

76. Röst, H. L. et al. TRIC: an automated alignment strategy for reproducible protein quantification in targeted proteomics. Nat Methods 13, 777–783 (2016).

77. Blattmann, P., Heusel, M. & Aebersold, R. SWATH2stats: An R/Bioconductor Package to Process and Convert Quantitative SWATH-MS Proteomics Data for Downstream Analysis Tools. PLoS One 11, e0153160 (2016).

78. Rosenberger, G., Ludwig, C., Röst, H. L., Aebersold, R. & Malmström, L. aLFQ: an R-package for estimating absolute protein quantities from label-free LC-MS/MS proteomics data. Bioinformatics 30, 2511–2513 (2014).

79. Choi, M. et al. MSstats: an R package for statistical analysis of quantitative mass spectrometry-based proteomic experiments. Bioinformatics 30, 2524–2526 (2014).

